# CROPseq-multi: a universal solution for multiplexed perturbation in high-content pooled CRISPR screens

**DOI:** 10.1101/2024.03.17.585235

**Authors:** Russell T. Walton, Yue Qin, Bryce Kirby, J. Owen Andrews, Mikko Taipale, Byunguk Kang, Paul C. Blainey

## Abstract

Forward genetic screens seek to dissect complex biological systems by systematically perturbing genetic elements and observing the resulting phenotypes. While standard screening methodologies introduce individual perturbations, multiplexing perturbations improves the performance of single-target screens and enables combinatorial screens for the study of genetic interactions. Current tools for multiplexing perturbations are limited by technical challenges and do not offer compatibility across diverse screening methodologies, including enrichment, single-cell sequencing, and optical pooled screens. Here, we report the development of CROPseq-multi (CSM), a CROPseq^1^-inspired lentiviral system to multiplex Streptococcus pyogenes (Sp) Cas9-based perturbations with versatile readout compatibility and high performance for both perturbation and barcode identification. CSM has equivalent per-guide activity to CROPseq and low lentiviral recombination frequencies. Dual-guide CSM libraries are constructed in a single, facile molecular cloning step that facilitates the use of unique molecular identifiers. CSM is compatible with enrichment screening methodologies, single-cell RNA-sequencing readouts, and optical pooled screens. For optical pooled screens, an optimized and multiplexed *in situ* detection protocol improves barcode counts 10-fold (for mRNA detection), enables detection of recombination events, and reduces the number of sequencing cycles required for decoding by 3-fold relative to CROPseq. CROPseq-multi-v2 (CSMv2) adds compatibility for detection methods based on T7 RNA polymerase *in vitro* transcription^2–5^. CSM provides a single system for CRISPR screens that is compatible with individual and combinatorial perturbations, diverse SpCas9-based perturbation technologies, and multiple high-content, single-cell phenotypic readouts.

## Introduction

Most pooled screens achieve perturbation of genetic components through lentiviral delivery of one or more Cas9 single guide RNAs (sgRNAs). Lentiviral delivery systems are near-ubiquitous across *in vitro* pooled screens conducted today due to their ability to efficiently deliver perturbations to a wide range of cell types, to control copy number via multiplicity of infection (MOI), and to retain a record of the perturbation identity integrated in the host-cell genome. Most commonly, a single perturbation is delivered to each cell, however multiplexing perturbations offers several advantages. Despite improvements in algorithms for guide design^6–11^, the selection of active and specific guides remains challenging and pairing guides can increase on-target performance. The use of multiple guides targeting the same genetic element (“single-target screens”) has been shown to improve on-target activity in CRISPR knockout (CRISPR-KO), activation (CRISPRa), and interference (CRISPRi) screens^12–15^. Multiplexing perturbations also enables targeting multiple distinct genetic elements per cell (“combinatorial screens”) to identify genetic interactions – phenotypes that arise from the interaction of genetic components.

Multiplexing perturbations in screens typically requires multi-perturbation lentiviral vector designs (“multiplexing vectors”). In principle, single-perturbation vectors can enable random sampling of perturbation combinations in pooled screens via either high MOI (with limited control of multiplicity) or serial transduction and selection^16,17^. However, random sampling of perturbation combinations requires profiling an immense combinatorial space, which scales as 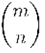, for *m* targets and combinations of *n* perturbations per cell. For example, pairwise interactions of only 200 gene targets results in 19,900 unique combinations, roughly equivalent in scale to single-target genome-wide screens in common practice today. This limitation precludes the use of single-perturbation vectors for multiplexed single-target screens and poses a major limitation to their utility for combinatorial screens. In contrast, multiplexing vectors are delivered with a single selection step at a low MOI. Multiplexing vectors uniquely enable multiplexing in single-target screens and offer an alternative to random sampling in combinatorial screens. In addition to enabling an exhaustive search of the combinatorial space, multiplexing vectors allow the perturbation of a biologically informed subset of target combinations^18–22^. In such screens, the size of multiplexed vector libraries scales linearly with the number of selected target combinations.

Existing lentiviral perturbation systems offer performance and functionality tradeoffs with respect to multiplexing capacity, lentiviral recombination frequencies, and barcode readout compatibility (**Supplementary Table 1**). Derived from standard single-perturbation designs (**Supplementary Figure 1A**), vectors encoding serial pol III (typically U6) promoters and SpCas9 sgRNAs (“guides”) are a common approach to multiplex perturbations^23,13,24^ (**Supplementary Figure 1B**). A major limitation of these multiplexing systems is the roughly 400 base pair (bp) distance separating sgRNA spacer sequences, leading to lentiviral recombination and unintended sgRNA combinations in about 30% of cells (**Table 1**)^23,13,24^. These recombination events are an inherent property of lentiviral systems as lentivirus are pseudodiploid and, in the process of infection, reverse transcription during minus-strand synthesis exhibits frequent template switching in a homology and distance-dependent manner^25–27^ (**Supplementary Figure 1C, D**). If perturbation identities are assigned based on the observation of only one sgRNA, these recombination events will be misassigned and contribute to experimental noise^25^. If the screening methodology captures both sgRNA identities, recombination events can be detected and filtered out. However, high recombination rates remain a major burden when perturbation coverage is challenging or costly to maintain, such as in single-cell RNA-sequencing (scRNA-seq)-based screens^24^.

**Table 1.** Comparison of selected lentiviral gRNA delivery systems for barcoding and multiplexing. *Up to 2 for pooled library construction; up to 3 have been demonstrated for arrayed library construction^23^. **Predicted based on distance. BC, barcode.

| <u>vector</u> | <u>Cas effector(s)</u> | <u>perturbation identifier</u> | <u>spacer-BC recombination</u> | <u>mRNA barcode</u> | <u>multiplex</u> | <u>spacer- spacer recombination</u> |
| --- | --- | --- | --- | --- | --- | --- |
| pLentiGuide and similar | SpCas9 | spacer | n/a | no | up to 2* | 26%<sup>68</sup> |
| LentiGuideBC and similar | SpCas9 | linked BC | 50%<sup>25</sup> | yes | no | n/a |
| CROPseq | SpCas9 | spacer | n/a | yes | no | n/a |
| Big Papi | SpCas9 and SaCas9 | linked BC | 5%<sup>30</sup> | no | 2 | 9%<sup>30</sup> |
| Cas12a crRNA array | Cas12a | spacer | n/a | no | 3+ | <1%** |
| **CROPseq-multi** | SpCas9 | spacer or linked BC | <1% (this work) | yes | up to 2 | 12% (this work) |

Other designs seek to minimize recombination by reducing the distance separating sgRNAs. The Big Papi vector^28^ and similar designs^29^ employ antiparallel orthogonal pol III promoters and Cas9 sgRNAs, reducing the distance separating spacers to under 200 bp and decreasing recombination to about 9%^30^. Other systems substitute Cas9 for Cas12a, capitalizing on the native crRNA array processing ability of Cas12a enzymes^31,32^, enabling separation of spacers by only 20 bp and likely reducing recombination to negligible levels^14,33,29,12^. However, in contrast to the widely used SpCas9, Cas12a enzymes are limited in guide design by relatively restrictive protospacer adjacent motifs, are less effective in CRISPR-KO screens unless compensated by multiple Cas12a guides per gene^14,29,33^, lag in development with applications including CRISPR-KO, CRISPRa, CRISPRi, base editing, and prime editing^34–36^, and are limited in the availability of existing cell lines and animal models. Regardless, all current multiplexing solutions lack compatibility with mRNA-barcoding, which is required for some screening modalities, including popular single-cell RNA-sequencing and *in situ* detection workflows^1,16,23,37^.

LentiGuide-Barcode (LentiGuideBC)^16^ vectors and similar designs (e.g. Perturb-seq^37^, MOSAIC-seq^38^, CRISP-seq^39^) typically sacrifice multiplexing capability in pooled screens for the ability to express a linked barcode in mRNA (**Supplementary Figure 1E**). In principle, these vector designs are not fundamentally incompatible with multiplexing; however, constructing libraries with at least three distal programmed sequence elements (i.e. two sgRNAs and a barcode) is challenging. To date, these designs have required multi-step arrayed cloning^23^, making the approach impractical for most pooled screens. In these designs, the linked barcode is separated from the sgRNA spacer by at least 1,700 bp, comprising the pol II promoter and selection gene, resulting in lentiviral recombination near the theoretical maximum of 50% and contributing to a major loss of statistical power^25^. Efforts to decrease this distance by moving the U6-sgRNA downstream of the pol II promoter and resistance gene have resulted in poor guide activity, potentially due to transcriptional interference^1,25^. Our group described co-packaging integration-deficient templates as a means to mitigate recombination, albeit at the cost of a 100-fold reduction in lentiviral titer^26^.

The CROPseq approach offers a solution to mRNA-barcoding that is not impacted by lentiviral recombination^1^ (**Supplementary Figure 2A, B**). By embedding the U6 promoter and sgRNA within the lentiviral 3’ long terminal repeat (LTR), the CROPseq design leverages the high-fidelity intramolecular duplication of the 3’ LTR to the 5’ end of the lentiviral genome during genome integration (**Supplementary Figure 2B**). This duplication results in two copies of the sgRNA, with the 5’ copy expressing functional sgRNAs without transcriptional interference and the 3’ copy transcribed as mRNA in the 3’ untranslated region (UTR) of the pol II-transcribed selection gene, compatible with mRNA-detection approaches. With these considerations in mind, we sought to engineer a lentiviral system to enable multiplexed SpCas9-based perturbation with mRNA-barcoding.

## Results

### Design of a CROPseq-inspired multiplexing vector

We reasoned that the lentiviral 3’ LTR could be compatible with a minimal multiplexing system. While CROPseq vectors enable faithful duplication of the 3’ LTR via intramolecular recombination during plus-strand synthesis, adjacent guides and/or barcode elements are still vulnerable to intermolecular recombination during minus-strand synthesis (**Supplementary Figure 2C**). To minimize the 3’ LTR insertion and sgRNA separation, we opted to use endogenous transfer RNA (tRNA) processing as our multiplexing solution, as others have implemented^40–46^ (**Figure 1A**). Encoded in about 72 bp, tRNA molecules recruit endogenous RNAses P and Z to cleave the tRNA at the 5’ and 3’ ends, such that, when positioned between sgRNAs within a single transcript, the RNA is processed into separate, functional sgRNAs. While only a single tRNA is required for multiplexing (i.e. U6-sgRNA-tRNA-sgRNA), we additionally preceded the first sgRNA with a tRNA (i.e. U6-tRNA-sgRNA-tRNA-sgRNA) as tRNAs encode their own promoter elements and, in some contexts, may improve expression alone or in conjunction with a U6 promoter^40,45,46^. Additionally, each tRNA eliminates the requirement for a 5’ guanine base on the following guide that is otherwise required in U6 transcription systems and is often encoded as a mismatched 20th or 21st base of the spacer. This design increased the size of the 3’ LTR insertion from 352 bp in CROPseq to 643 bp in CROPseq-multi (**Supplementary Figure 3A**). tRNA-encoding sequences can be processed by the endogenous RNases out of mRNA^46^ and processing of tRNAs out of the lentiviral genomic RNA and the selection gene transcript would interfere with lentiviral production, selection, and detection of barcoded mRNA. To enable selective processing of tRNAs from pol III (but not pol II) transcription products, we reversed the orientation of the elements within the 3’ LTR, such that the U6 promoter, tRNAs, and sgRNAs are encoded on the lentiviral minus strand. We termed our CROPseq-inspired multiplexing solution **CROPseq-multi (CSM)**.

**Figure 1.**
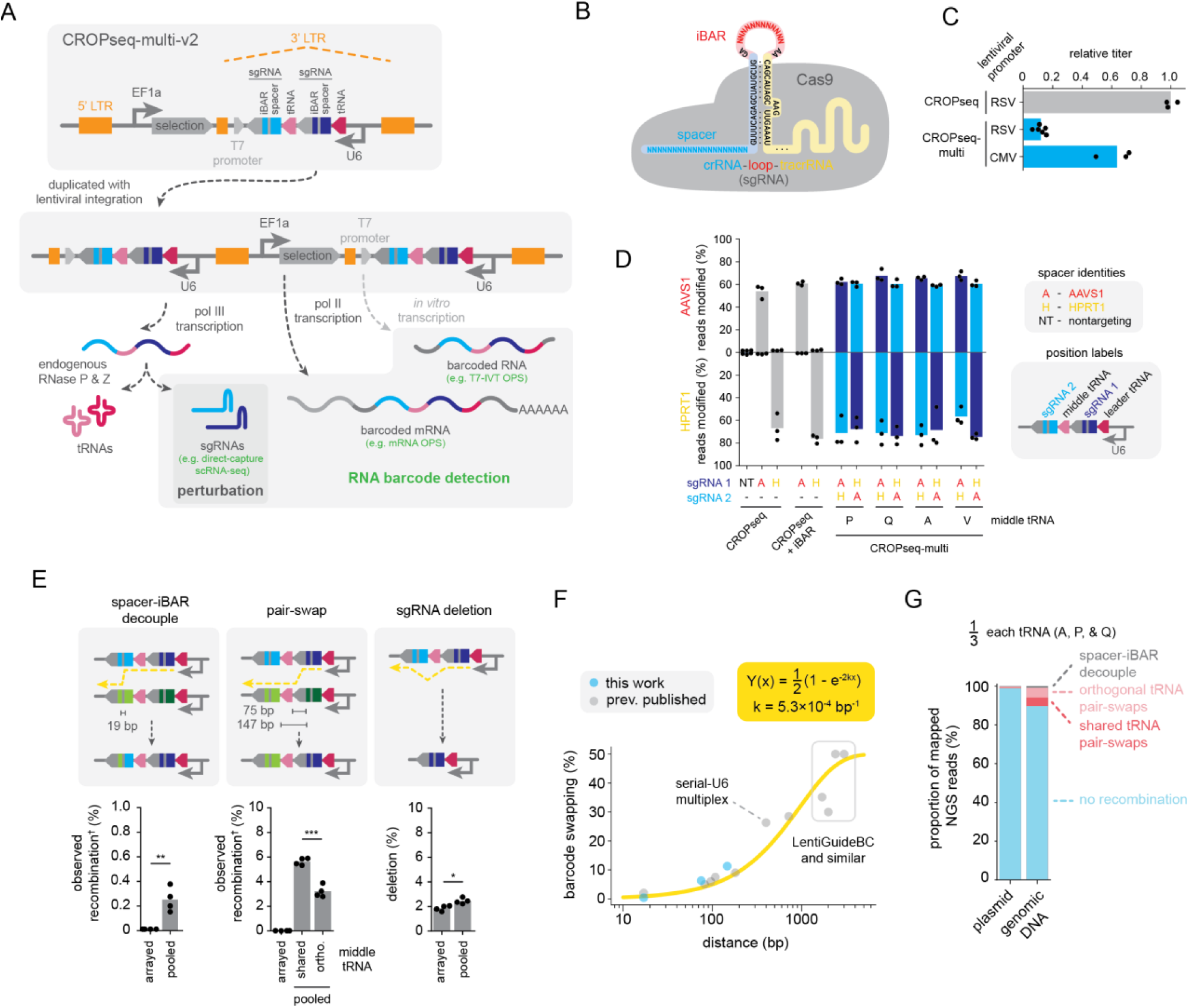
CROPseq-multi: a versatile sgRNA multiplexing system. (A) CROPseq-multi encodes two sgRNAs with internal barcodes (iBARs), multiplexed using tRNAs, within the lentiviral 3’ long terminal repeat (LTR). The 3’ LTR is duplicated during lentiviral integration, producing a second copy of the sgRNAs. (B) iBARs place linked barcodes within the loop joining the crRNA and tracrRNA into the sgRNA. (C) Functional lentiviral titers of CROPseq-multi relative to CROPseq. (D) Genome editing activities of CROPseq and CROPseq-multi vectors in SpCas9-expressing A549 lung adenocarcinoma cells, quantified by next-generation sequencing. Mean and n=3 biological replicates shown. (E) Observed lentiviral recombination of CROPseq-multi barcode elements. (T-test: *p<0.05, **p<0.01, ***p<0.001) Illustrations depict one example of each type of recombination. ^†^In this assay, observed recombination is expected to reflect half the total recombination (see Supplementary Figure 3D). (F) Lentiviral barcode swapping rates as a function of the distance separating barcode elements (e.g. spacer-spacer, spacer-linked barcode, etc.) in published barcoding systems and CROPseq-multi (see Supplementary Table 1). (G) Recombination in a CROPseq-multi library with equal proportions of three orthogonal tRNAs in plasmid DNA (prior to lentiviral recombination) and genomic DNA (after lentiviral recombination). scRNA-seq, single-cell RNA-sequencing; OPS, optical pooled screens; IVT, *in vitro* transcription; LTR, long terminal repeats; RSV, Rous sarcoma virus; CMV, human cytomegalovirus.

To add compatibility with approaches that use *in vitro* transcription with T7 RNA polymerase (T7-IVT) for detection of perturbation barcodes^4,5^, we included a T7 promoter in the 3’ LTR in an updated design, termed **CROPseq-multi-v2 (CSMv2)** (**Figure 1A, Supplementary Figure 3A**). Throughout, for applications requiring T7-IVT compatibility, we refer specifically to **CSMv2**, otherwise we refer to both generically as **CSM** as they are otherwise interchangeable.

In addition to our guide multiplexing changes, we added 12 bp barcodes internal to the sgRNAs (iBARs) as freely specified additional readout elements linked to the sgRNA^47^ (**Figure 1A, B, Supplementary Figure 3A**). Linked barcodes are advantageous to minimize the sequence length needed to uniquely identify library members, to represent combinations of guides that may not be individually unique, and to encode additional information such as clonal identity. As linked barcodes are prone to distance-dependent recombination, the iBAR system is attractive for the placement of barcodes only 19 bp from the spacer, within the synthetic loop that joins the crRNA and tracrRNA into a sgRNA (**Figure 1B, Supplementary Figure 3A**). As the iBARs are transcribed both antisense as mRNA and within the sgRNA scaffold, their detection is compatible with both mRNA-based^16,23,37,39^ and direct-capture^13^ (pol III product) protocols.

We first addressed the compatibility of CSM with lentiviral production and transduction. It has been suggested that the lentiviral 3’ LTR may be incompatible with the larger insertions needed to encode multiple perturbations^25,27^. Insertions of up to 1,200 bp within the 3’ LTR have been evaluated, albeit with reduced viral titers^48^. While our initial design was compatible with lentiviral delivery, we observed a roughly 10-fold reduction in functional titer compared to CROPseq (**Figure 1C**). Reasoning that increased size of the 3’ LTR insertion might explain the low titer, we compared viral titers of CSM with one guide (445 bp insertion) or two guides (643 bp insertion) against CROPseq (352 bp insertion). Surprisingly, we observed the same 10-fold reduction in titer with the single-guide CSM, suggesting that titer did not decrease linearly with increasing 3’ LTR insertion size and that the sequence elements (i.e. tRNA(s)) and/or orientation may contribute (**Supplementary Figure 3B**). We hypothesized that an alternate lentiviral promoter could improve titer if transcriptional interference with the U6 promoter and tRNAs was hindering lentiviral genome production. While most CROPseq vectors employ a Rous sarcoma virus (RSV) promoter to drive lentiviral genome expression during viral production, the human cytomegalovirus (CMV) promoter can improve titer in some lentiviral vectors^49^. Swapping the RSV promoter with a CMV promoter rescued the titer of the full two-guide CSM system to 64% that of CROPseq (**Figure 1C**). We proceeded to use the CMV promoter as the default construction for all CSM vectors.

### Genome editing performance of CSM

To assay the performance of CSM for genetic perturbation, we transduced SpCas9-expressing A549 lung adenocarcinoma cells with CROPseq and CSM vectors encoding guides targeting the AAVS1 and HPRT1 loci and evaluated genome editing efficiency by next-generation sequencing (NGS) (**Figure 1D**). Ideally, multiplexing systems possess equivalent per-guide activity to single-plex systems and display minimal positional bias (i.e. equal guide activity in all positions). We first validated that 12 bp iBARs were not detrimental to activity in the CROPseq vector architecture (**Figure 1D**). Next we tested both orientations of spacers and four guide-intervening “middle” tRNAs in CSM. None of the tested CSM constructs had significantly different activity for either target compared to CROPseq or one-another (T-test with Bonferroni correction) (**Figure 1D**). We selected designs employing human proline, glutamine, and alanine tRNAs (tRNA_P_, tRNA_Q_, and tRNA_A_, respectively; see **Supplementary Table 2**) for further development because they displayed the most consistent activity of the four tRNAs tested. CSM and CSMv2 were indistinguishable with respect to genome editing performance (**Supplementary Figure 3C**).

### Lentiviral recombination with CSM

To evaluate lentiviral recombination rates, we performed arrayed and pooled lentiviral production with combinations of CSM vectors and measured the identities of barcode elements after transduction and integration in genomic DNA. Arrayed lentiviral preparation guarantees that co-packaged genomes are identical, meaning that template switching does not disrupt perturbation pairing, and serves as a control for other sources of recombination, such as polymerase chain reaction (PCR) amplification. In pooled lentiviral preparation, as is performed in pooled screens, lentiviral genomes are essentially randomly paired within virions. In a pooled preparation of two vectors, 50% of virions will harbor different perturbation pairs that can result in observable intermolecular recombination events (**Supplementary Figure 3D**). In a high-complexity library, nearly all virions are expected to package different perturbation pairs, so the recombination rate should approach 2x the observed rate in a two-vector assay. In all conditions, recombination was measured by NGS of genomic integrations.

Intermolecular recombination events that decoupled spacers from their iBARs were detected at rates below 1%, likely acceptable for most applications (**Figure 1E**). For two CSM vectors with a common middle tRNA, intermolecular recombination events resulting in incorrect pairings of sgRNAs (“pair-swaps”) were observed at about 5.9% (**Figure 1E**). As the middle tRNAs we validated are divergent in sequence (**Supplementary Figure 3E**), we reasoned that if co-packaged CSM constructs encoded orthogonal middle tRNAs, the homologous barcode-separating distance would be reduced to only 75 bp (**Supplementary Figure 3A**). In a pooled screening setting, this would be implemented by designing libraries with multiple orthogonal middle tRNAs to reduce the probability of co-packaging constructs with a common middle tRNA. Correspondingly, the use of orthogonal tRNAs reduced observed pair-swap recombination events to 3.2% (**Figure 1E**). We converted observed rates to expected underlying rates and compared with previously reported recombination frequencies from other dual barcode systems (either multiplexed guides or guides with linked barcodes) (**Figure 1F, Supplementary Table 1**). Modeling barcode swapping frequency as a function of the length of homologous intervening sequence, we found that lentiviral recombination frequency could be modeled by fixed per-bp probability of template switching of 5.3x10^-4^ per bp, within the range of previously reported measurements of 4.9-13.5x10^-4^ per bp^50^ (**Figure 1F**).

With both arrayed and pooled lentiviral preparations, we observed sgRNA deletion events in about 2% of integrations that appeared to be driven by sgRNA scaffold homology (**Figure 1E**), despite the use of two orthogonal scaffold sequences (**Supplementary Figure 3F**). These events exclusively result in the deletion of the second spacer, an observation that is not readily explained by chimeric PCR artifacts, but is consistent with lentiviral recombination given the unidirectionality of lentiviral reverse transcription (**Figure 1E, Supplementary Figure 1C**). The resulting guide scaffolds are chimeras of the two orthogonal scaffolds and may vary in functionality. Examination of recombination outcomes revealed that the modest but significant increase in deletion frequencies in the pooled condition was driven by spacer homology – likely an artifact of the use of identical spacers in opposite positions in two of the three constructs used in our assay (i.e. sgRNA_AAVS1_-tRNA-sgRNA_HPRT1_ and sgRNA_HPRT1_-tRNA-sgRNA_AAVS1_) – and should not be a factor in standard high-complexity libraries. Considering the observed deletion rates and the expected underlying spacer-iBAR decoupling and pair-swap rates based on the observed rates, we estimated the fraction of CSM integration events without recombination for complex libraries to be 86% with the use of a single middle tRNA or 90% when implemented with three orthogonal middle tRNAs (**Supplementary Figure 3G**). We opted to use three orthogonal middle tRNAs in equal representation for all complex libraries moving forward.

### Construction and quality control of pooled CSM libraries

We next validated the construction of pooled CSM libraries. We first evaluated a one-step assembly protocol (**Supplementary Figure 4A**) for sequence fidelity, recombination, and uniformity for a 1,080-member library encoding three different middle tRNAs in equal representation. Mapping rates of individual barcode elements (spacer 1, iBAR 1, middle tRNA, spacer 2, and iBAR 2) ranged from 94-98% and all elements together mapped jointly to the library as designed in 84% of reads, with NGS error rates likely contributing to a substantial fraction of unmapped reads (**Supplementary Figure 4B**). We initially found that libraries were prone to PCR-mediated recombination during oligo pool amplification. Optimization of the amplification conditions revealed that PCR-mediated recombination was almost entirely determined by oligo pool template concentrations and the use of low template concentrations (12 pg/µL) could effectively eliminate recombination (**Supplementary Figure 4C**). With optimized amplification conditions, we observed less than 1% recombination in our plasmid libraries (**Figure 1G, Supplementary Figure 4C**). This oligo pool amplification strategy did not compromise library uniformity when scaled to the appropriate number of amplification reactions (see **Methods**) and, when paired with an optimized assembly procedure, resulted in 90:10 ratios (the ratio of the 90th and 10th percentiles of abundance) below 2 and Gini coefficients below 0.14 (**Supplementary Figure 4D-F**). Construction of additional libraries with these optimized conditions, ranging in size from about 100 to 40,000 members, yielded reproducible mapping (90% mean ± 5% standard deviation for spacers), uniformity metrics (90:10 ratios of 2.2 mean ± 0.5 standard deviation and Gini coefficients 0.17 mean ± 0.05 standard deviation), and recombination (0.8% mean ± 0.6% standard deviation) (**Supplementary Figure 4G**).

We also optimized an alternate oligo pool amplification strategy to repurpose the second iBAR sequence as a clonal barcode (**Supplementary Figure 4H**). This approach requires no changes to oligo pool design, only alternate amplification primers, to generate libraries with high complexity clonal barcodes (**Supplementary Figure 4H, I**).

Following pooled lentiviral preparation and transduction, we assayed recombination in genomic DNA and observed 90.5% of genomic integrations without recombination (**Figure 1G**). Across eight plasmid libraries and four cell lines, we observed a range of 9-17% total (plasmid + lentiviral) recombination, with 8-15% (mean 12%) attributable to lentiviral integration (**Supplementary Figure 4J**). This analysis excludes sgRNA deletion events that were not captured by a modified NGS library preparation procedure but otherwise recapitulates the expected contributions to recombination from spacer-iBAR decoupling and pair-swap events based on our arrayed measurements (**Figure 1D, Supplementary Figure 3G**).

### Pooled viability screens validate the performance of CSM for CRISPR-KO and CRISPRi and single-target and combinatorial screens

We performed a series of viability screens to evaluate the performance of CSM in a common pooled screening task (**Figure 2A**). To assay performance in diverse settings, we used three model systems and two perturbation modalities (**Figure 2B**). For CRISPR-KO screens, we used HeLa cells with doxycycline-inducible expression of Cas9 as well as wild-type A549 cells in which Cas9 was delivered “all-in-one”, on the CSM vector itself. For the CRISPRi screen, we used RPE1 cells with constitutive expression of dCas9-KRAB^24^.

**Figure 2.**
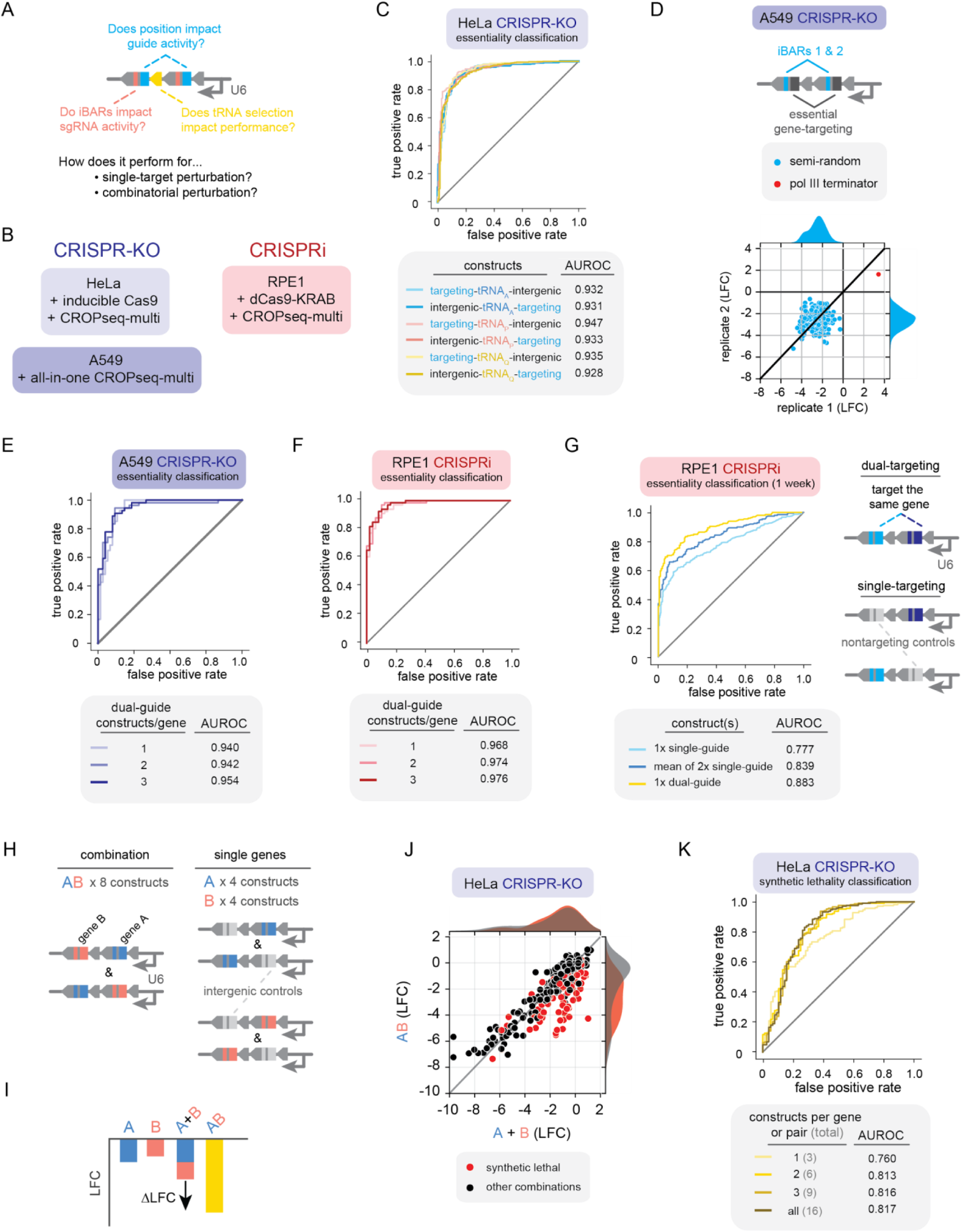
Benchmarking CROPseq-multi in pooled viability screens. (A) Performance metrics addressed with pooled viability screening. (B) Cell line and perturbation approaches employed. (C) Gene essentiality classification varying the gene targeting guide position and middle tRNA identity. (D) Variability in the performance of constructs differing by only 12 nt iBAR sequences. (E-F) Gene essentiality classification with one to three *a priori* ranked dual-guide constructs per gene, where each construct pairs two guides targeting the same gene, for CRISPR-KO (E) and CRIPSRi (F) screens. (G) Gene essentiality classification for individual dual-guide constructs, individual single-guide constructs, and the average of two single-guide constructs. (H) The design of constructs in a combinatorial perturbation library targeting individual genes (A or B) and combinations (AB). (I) Schematic representation of a synthetic lethal genetic interaction, quantified with log2 fold changes in viability screening. (J) Depletion of gene pairs in combination (AB) versus the expected depletion (A + B) based on the depletion of individual genes (A and B). Gene pairs in red are previously reported synthetic lethal pairs. (K) Classification of synthetic lethal gene pairs for variable numbers of constructs per gene and gene pair. For all panels, data is from a two-week viability timepoint unless stated otherwise. LFC, log2 fold change; AUROC, area under receiver operator characteristic curve.

To test for any systematic bias in guide position and tRNA performance, we compared 3,444 constructs pairing intergenic control guides with guides targeting genes of variable essentiality, alternating the order of the guides and middle tRNA identity. We assayed representation after a two-week viability screen in HeLa cells and used log2 fold change (LFC) values to classify gene essentiality. All combinations of guide order and middle tRNA performed comparably with area under receiver operator characteristic curve (AUROC) values between 0.93 and 0.95 (**Figure 2C)**. We further tested 342 unique gene-targeting guide pairs (both guides targeting the same gene) in a two-week viability A549 screen, targeting 114 genes of variable essentiality, in both orientations and with all three tRNAs. Again we observed concordant performance among middle tRNAs for gene essentiality classification (AUROC 0.912-0.924 for individual constructs) (**Supplementary Figure 5A**). The variability between individual constructs harboring opposite guide orientations was comparable to the variability between biological replicates (**Supplementary Figure 5B**).

Additionally, to evaluate the impact of iBAR sequences on guide activity, we selected guide pairs to target essential genes DBR1, PCNA, and RPA3 and compared 200 constructs per target varying only by 12 nt iBAR sequences. Essentiality classification was nearly perfect for all targets (AUROC 0.995-0.998) and there was little to no variability in performance of constructs with different iBARs beyond the biological replicate variability (**Figure 2D**, **Supplementary Figure 5C**).

Previous studies have demonstrated that multiplexing systems offer improved performance over single-guide systems for single-target screens, enabling the use of smaller libraries. We performed a CRISPRi screen in RPE1 cells and a CRISPR-KO screen in A549 cells, targeting the same 114 genes with a range of essentiality scores in DepMap^51,52^, with three dual-guide constructs per gene (with both guides targeting the same gene). We ranked constructs *a priori* based on computational guide rankings generated with the guide design tool CRISPick^8,9^. For CRISPR-KO, we designed constructs pairing the top two ranked guides, the third and fourth ranked guides, and the fifth and sixth ranked guides. For CRISPRi, we similarly paired the top two guides together, the third and fourth guides together, and then evaluated a guide pair from a published screen for the third pair^24^.

For CRISPR-KO, we were able to classify essential genes with an AUROC of 0.940 with just one construct per gene, or 0.954 using the mean of all three constructs (**Figure 2E**). For CRISPRi, essentiality classification performed similarly with an AUROC of 0.968 with just one construct per gene, or 0.976 for three constructs (**Figure 2F**). For both CRISPR-KO and CRISPRi, we observed that the relative performance of constructs was consistent with the predicted guide rankings (**Supplementary Figure 5D, E**).

For the CRISPRi screen, we additionally decomposed each dual-guide construct into two corresponding single-guide constructs, each paired with a nontargeting control guide, to directly compare single-guide and dual-guide performance. For essentiality classification, dual-guide constructs (AUROC 0.940) outperformed both the individual guides (AUROC 0.868) and composite metrics for corresponding pairs of single guides (mean LFC, AUROC 0.928) (**Supplementary Figure 5F, G)**. The difference in performance was even more pronounced at an earlier timepoint, with dual-guide constructs (AUROC 0.883) outperforming individual guides (AUROC 0.777) and pairs of single-guides constructs (mean LFC, AUROC 0.839) (**Figure 2G, Supplementary Figure 5H**). Dual-targeting constructs had an average depletion more than 2-fold greater in magnitude compared to single-targeting constructs for essential genes (**Supplementary Figure 5I**). Consistent with previous studies^12,13,15^, these results suggest that multiplexed perturbation systems not only outperform single-perturbation systems for single-target screens, but can do so with smaller library sizes.

Guide multiplexing systems are also critical to enable perturbation of programmed combinations of genes for the study of genetic interactions. We performed a CRISPR-KO screen in HeLa cells, targeting 280 gene pairs including 76 pairs previously identified as synthetic lethal HeLa cells^53^ or among 13 “gold standard” pairs that exhibit synthetic lethality across multiple screening datasets^54^ and 83 pairs reported to have no viability interaction^53^. We designed 4 constructs for each individual gene (A and B), pairing the gene targeting guide with an intergenic control guide, and eight constructs targeting both genes in combination (AB) (**Figure 2H**). In viability screens, a gene pair is defined as synthetic lethal if the viability defect of the combination exceeds the additive effects of the individual gene perturbations and can be quantified by the difference in LFC (ΔLFC) between the combination (AB) and the sum of the individual perturbations (A+B) (**Figure 2I**). We observed a strong deviation from additive effects for many reported synthetic lethal pairs (**Figure 2J).** Though interactions commonly have low reproducibility rates across screens^54^, we evaluated our ability to identify interactions previously reported in one published screen in HeLa cells^53^. We were able to classify synthetic lethality from ΔLFC scores with an AUROC of 0.817 (**Figure 2K**). Downsampling the number of constructs revealed only modest improvement beyond the top two *a priori* ranked constructs per gene or gene pair (six constructs in total for A, B and AB), with an AUROC of 0.813 (**Figure 2K**). This design suggestion – 6 constructs per gene pair – matches the recommendations for current state-of-the-art Cas12 systems with 4 guides per construct^54^.

Having benchmarked the performance of CSM in single-target and combinatorial pooled viability screens, we next focused on applicability for high-content screens, in particular single-cell RNA-sequencing screens and optical pooled screens.

### CSM is compatible with scRNA-seq screens

Screens with a scRNA-seq readout (i.e. CROP-seq^1^, Perturb-seq^23,37^, CRISP-seq^39^, etc.) provide rich, single-cell-resolved transcriptional measurements paired to genetic perturbations. However, implementations for both individual^37,55^ and multiplexed^23,24^ perturbations have been hindered by challenges in barcode detection and lentiviral recombination^25,27^. CROPseq and CSM vectors produce both pol II (mRNA) and pol III (sgRNA) transcription products containing perturbation-identifiers (i.e. spacer, iBAR), providing multiple options for barcode detection.

We chose to evaluate barcode capture with a droplet-based 5’ capture scRNA-seq workflow that detects pol III product sgRNAs, also known as direct-capture^13^. The minor modifications required for 5’ capture scRNA-seq with CSM are the addition of two reverse transcription primers, one for each guide, and a single multiplexed PCR to enrich sgRNAs from the whole transcriptome amplification product (**Figure 3A**). We selected this workflow because of the wide adoption and commercial availability, ubiquity and functional relevance of pol III expression, compatibility with V(D)J capture, and demonstrated performance advantages^13^. With appropriate modifications to enrich for barcoded transcripts analogous to published methods^13,56,57^, we anticipate that CSM should be compatible with a diversity of approaches that rely on capturing barcodes in either pol II or pol III products. CSMv2 will likely offer an advantage over the original CSM for strategies that use T7-IVT for barcode enrichment^57^.

**Figure 3.**
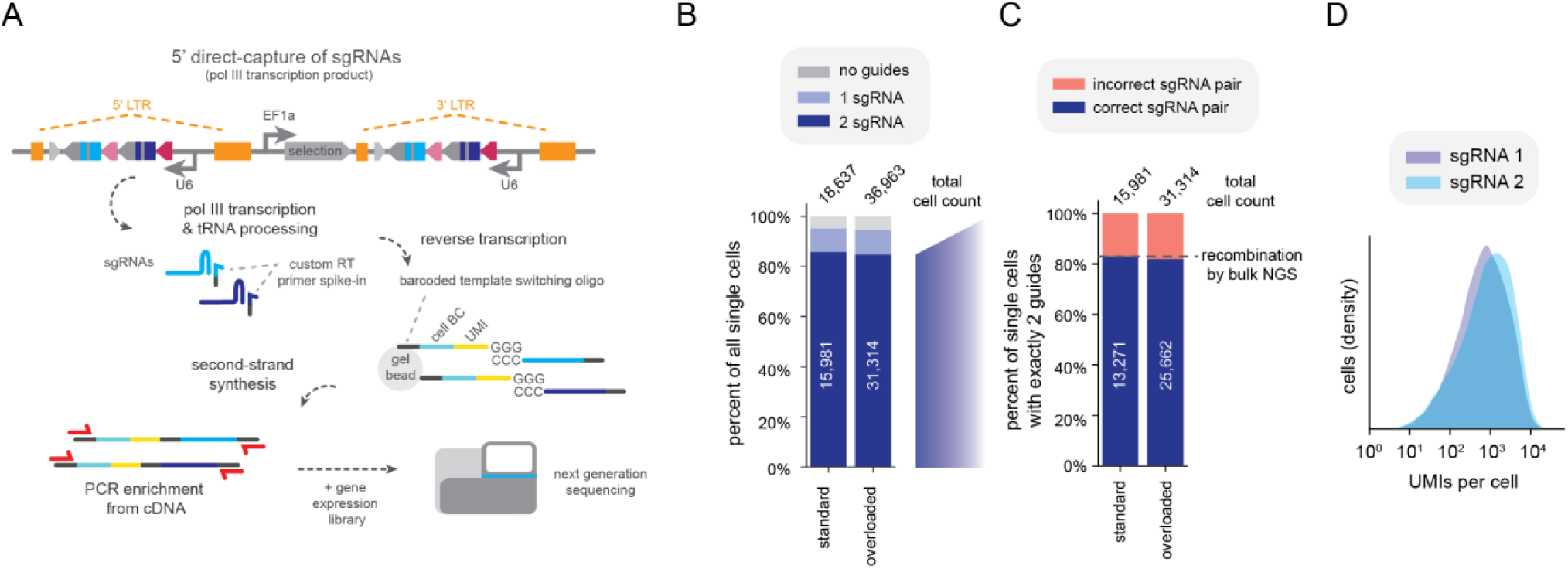
CROPseq-multi for single-cell RNA-sequencing screens. (A) Schematic representation of 5’ direct-capture (pol III transcription products) for a single-cell RNA-sequencing readout with CROPseq-multi. (B) Percentage of all single cells (singlet events) by the number of detected sgRNAs in HeLa cells. (C) Percentage of cells with exactly 2 sgRNAs detected with correct and incorrect sgRNA pairs. (D) Unique molecular identifiers per sgRNA per cell for cells with exactly 2 sgRNAs detected. LTR, long terminal repeats; RT, reverse transcription; cell BC, cell barcode; UMI, unique molecular identifier; NGS, next generation sequencing.

We evaluated barcode capture in HeLa cells with both standard cell loading (∼29,000 cells per GEM-X lane) and overloading (∼83,000 cells per GEM-X lane), which enables increased cell recovery at the cost of higher multiplet rates. Standard and overloading yielded a total of 21,973 and 51,035 cells, respectively, before filtering multiplets. With robust barcode detection, perturbation barcodes can be used to filter multiplets, analogous to cell hashing approaches^58^. Under both loading conditions, only 4% of cells had no guides detected and the most frequent occurrences were two unique guides per cell, i.e. singlets, followed by four unique guides per cell, i.e. doublets (**Supplementary Figure 6A, B**). We observed 16% and 27% of cells with more than two guides detected for standard and overloading conditions, respectively, consistent with the expected increase in multiplets with overloading (**Supplementary Figure 6C**). An increase in the total number of reads and genes per cell in cells with more than 2 guides, compared to those with 0-2 guides, further supports that these events reflect multiplets (**Supplementary Figure 6D-G**). Excluding multiplets left 18,637 cells with standard loading and 36,963 cells with overloading.

After excluding multiplets, both loading conditions yielded 84.7-85.7% of cells with two guides detected, 9.4-9.8% of cells with one guide detected, and 4.8-5.5% of cells without guides (**Figure 3B**). Correct guide pairs comprised 82-83% of cells, closely recapitulating the 17% recombination rate in this cell library as measured by bulk NGS (**Figure 3C**). Overall, from standard and overloading conditions, we recovered 13,271 and 25,662 cells, respectively, in which exactly two guides reflecting a correct pair were observed. In cells with exactly two guides detected, we observed similar UMI counts for both guide positions under standard (**Figure 3D**) and overloading (**Supplementary Figure 6H**) conditions. Cells with only one guide detected were missing sgRNA 1 or sgRNA 2 with similar frequencies (**Supplementary Figure 6I**), ruling out guide deletion recombination events, which selectively delete spacer 2, as the primary explanation. Though recombination cannot be filtered out in these cells, their inclusion in downstream analysis may be acceptable in some contexts given low recombination rates. Of note, iBARs enable unique mapping of guide pairs even if only one guide is observed and spacer sequences are not unique.

### CSM enables sensitive detection and efficient decoding for optical pooled screens

Optical pooled screens enable high-content, single-cell-resolved phenotypic measurements, spanning spatial scales and including temporal dynamics and diverse molecular measurements, at the scale of tens of millions of cells^16,59,60^. However, the unique constraints of *in situ* barcode detection have made it challenging to multiplex perturbations with optical pooled screens. Briefly, *in situ* detection of barcodes for optical pooled screens involves fixation and permeabilization of cells, reverse transcription of barcoded mRNA to cDNA, fixation of the cDNA, copying the barcode sequence into a padlock probe (“gapfill”), ligation of the padlock into a circular ssDNA template, rolling circle amplification, and sequencing by synthesis to decode perturbations^16,61^ (**Figure 4A**). Robust mRNA expression is required for efficient detection and multiple perturbations must either be encoded by a single barcode or detected separately, as multiple barcodes. As sequencing reagent costs and imaging time impact screening throughput, minimizing the requisite number of sequencing cycles (i.e. barcode bases) to uniquely identify perturbations is desirable. Recently, approaches leveraging T7-IVT as an alternative to mRNA expression for optical pooled screens have also been described^4,5^. We validated the use of both mRNA and T7-IVT approaches for *in situ* detection with CSMv2.

**Figure 4.**
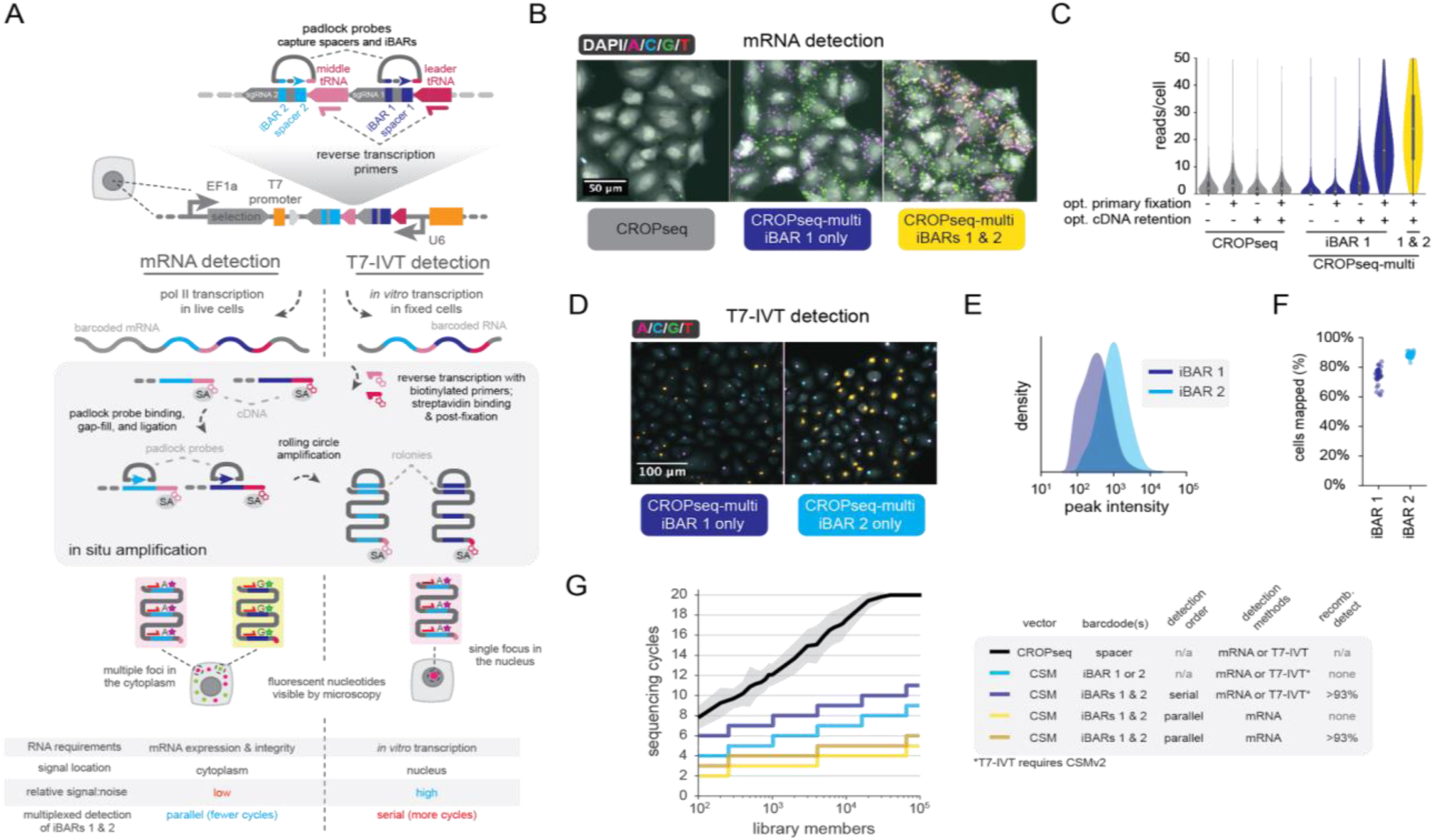
CROPseq-multi for optical pooled screens. (A) Overview of mRNA and T7 *in vitro* transcription (T7-IVT) *in situ* detection workflows. (B) Representative mRNA *in situ* sequencing images in A549 cells for CROPseq detection with the standard protocol and CROPseq-multi with an optimized protocol. (C) Optimizing *in situ* mRNA detection of CROPseq-multi. (D) Representative T7-IVT detection images of CROPseq-multi iBARs 1 and 2. (E-F) Quantification of peak intensity (E) and fraction of cells mapped (F) for T7-IVT detection protocols for iBARs 1 and 2. At least three technical replicates and three fields of view per technical replicate shown. (G) Modeling sequencing cycle requirements to uniquely identify library members with different decoding methods. For decoding via the spacer, libraries were simulated by randomly sampling guides from a genome-wide CRISPRi library (see Methods); mean and standard deviation are shown. CSM, CROPseq-multi; IVT, *in vitro* transcription; PFA, paraformaldehyde; SA, streptavidin; NGS, next-generation sequencing.

We initially optimized *in situ* detection in the context of mRNA-detection protocols. We designed padlock probes flanking each spacer and iBAR such that both are captured within the gapfill (**Figure 4A, Supplementary Figure 7A**). In contrast to the CROPseq vector, the orientation of *in situ* sequencing for CSM is opposite the orientation of the sgRNA (**Supplementary Figure 7B**). We first optimized the detection efficiency of the first spacer and iBAR pair in A549 cells transduced at an MOI of 0.1 with a library of three CSM vectors encoding orthogonal middle tRNAs. With our standard protocol, detection efficiencies (reads per cell) were low for CSM iBAR 1, averaging 1.3 reads per cell, compared to an average of 2.9 reads per cell for the CROPseq spacer (**Figure 4B, C**). We implemented two protocol changes to improve detection efficiencies for CSM (**Figure 4A**). First, we aimed to improve mRNA retention by altering the primary fixation, adding 0.007% glutaraldehyde to the standard 4% paraformaldehyde (PFA) fixative. Second, we sought to optimize cDNA retention by using a biotinylated reverse transcription primer and adding a streptavidin incubation between the reverse transcription and cDNA fixation steps to improve cDNA anchorage within the fixed cells (**Figure 4A**). The optimized cDNA retention alone improved detection efficiency to an average 7.5 reads per cell for CSM (**Figure 4B, C**). The optimized primary fixation alone did not substantially improve detection, but in combination with the optimized cDNA retention, detection efficiency improved further to an average 18.8 reads per cell (**Figure 4B, C**). These modifications had relatively modest effects on detection of the CROPseq vector (**Figure 4B**). In RPE1 cells, these protocol changes similarly improved detection efficiencies (**Supplementary Figure 8A, B**) and fine tuning of the primary fixation conditions suggested an optimal concentration of glutaraldehyde in the range of 0.007-0.003% in 4% PFA (**Supplementary Figure 8C**). Not all phenotypic measurements tolerate the addition of glutaraldehyde to the primary fixation step and it may be excluded at the cost of reduced detection efficiency (**Figure 4B, C, Supplementary Figure 8C**). CSM and CSMv2 perform identically for mRNA detection protocols (**Supplementary Figure 8D**).

An additional feature of the CSM design is the use of two barcodes to facilitate the detection of recombination events and to improve perturbation decoding efficiency. First, detection of both barcodes would enable identification and filtering of lentiviral recombination events. Second, with simultaneous detection of both barcodes as separate reads (“multiplexed detection”), a total of two nucleotides of a barcode pair can be decoded per sequencing cycle – one nucleotide from each barcode per cycle. Of note, this strategy is dependent on the ability to reliably detect both barcodes in each cell, which should be facilitated by the optimized detection protocol.

We evaluated multiplexed detection of iBARs 1 and 2 with our optimized protocol, using additional reverse transcription primers and padlock probes to detect the second spacer and iBAR pair. As these oligonucleotide reagents hybridize to sgRNA-adjacent tRNA, each orthogonal tRNA requires a different set of *in situ* detection oligos (**Supplementary Figure 7A**). For multiplexed detection with three orthogonal tRNAs, a total of four reverse transcription primers and four padlock probes are used, one set for the first spacer and iBAR and three sets (one for each middle tRNA) for the second spacer and iBAR. With multiplexed detection, we observed a mean of 27.7 total reads per cell in A549 cells (**Figure 4B, C**). Detection efficiencies were similar across constructs encoding different middle tRNAs (**Supplementary Figure 8E**). Multiplexed detection could impact per-barcode detection efficiencies due to optical crowding at high read densities. We evaluated detection of iBAR 1 and iBAR 2 both individually and multiplexed and observed only modestly lower per-barcode detection efficiencies when multiplexed (**Supplementary Figure 8F, G**). The absence of substantial spatial overlap of *in situ* sequencing reads may suggest that detection events for each spacer/iBAR largely originate from distinct mRNA molecules. We found that reads could be assigned to iBAR 1 or iBAR 2 based on either unique sequence mapping or the use of a fluorophore-conjugated oligo to label one of the iBARs (**Supplementary Figure 8H, I**). We used multiplexed *in situ* detection to decode perturbations and quantify recombination in cells transduced with three CSM constructs at an MOI of 0.1, prepared in either arrayed or pooled lentiviral settings. Assaying recombination with three unique vectors, the observed recombination rate is expected to reflect ⅔ of the underlying recombination rate, analogous to a two-vector recombination assay (**Supplementary Figure 3D**).

NGS of samples transduced with arrayed and pooled lentiviral preparations revealed observed pair-swap rates of <0.01% and 4.2%, respectively. Unlike NGS measurements, quantification of recombination *in situ* is sensitive to cell segmentation accuracy, technical artifacts such as transcript diffusion, and lentiviral MOI. We varied the stringency of read assignment to cells by varying the required minimum read counts per iBAR between 1 and 20, reasoning that cells with few reads for either iBAR might be the result of deletion recombination events, silencing of the lentiviral transgene, or incomplete selection, and, together with imperfect cell segmentation, could appear as false-positive pair-swap events (**Supplementary Figure 8H**). Relative to NGS, we observed modestly higher pair swap frequencies for the same samples *in situ*, ranging from 0.2-3.5% and 7.2-10.5% with arrayed and pooled lentiviral preparations, respectively, with the highest read count stringencies resulting in the lowest observed recombination frequencies (**Supplementary Figure 8H**). In a screening context, filtering out all such low-confidence assignments and incorrect pairings will be the primary objective so the distinction between pair-swap events and these potential modes of false-positives may be less important.

Next, we evaluated the use of T7-IVT-based approaches for optical pooled screens^4,5^ with CSMv2 (**Figure 4A**). We observed efficient detection of iBARs with both PFA fixation and decrosslinking (i.e. PerturbView^4^) (**Figure 4D**) and methanol/acetic acid fixation (i.e. NIS-seq^5^) protocols (**Supplementary Figure 9A**). With T7-IVT, detection of the T7-promoter-proximal barcode, iBAR 2, was more efficient than iBAR 1, consistent with the hypothesis that T7-IVT detection performance is sensitive to the promoter-barcode distance^4^. We observed brighter signals for iBAR2 (**Figure 4E**) and were able to map 88% of cells, compared to 74% for iBAR1 (**Figure 4F**). Despite the inter-iBAR variability, detection efficiencies for both iBARs remain comparable to published T7-IVT methods^4,5^. To reduce cost, we lowered the concentration of T7 RNA polymerase 5-fold while maintaining signal intensity and mapping rates (**Supplementary Figure 9B, C**).

With T7-IVT, the generation of just one primary *in situ* sequencing read per lentiviral integration complicates a straightforward application of multiplexed decoding, which relies on spatially distinct reads for iBARs 1 and 2. Instead, we prioritized a serial decoding scheme. First, one iBAR can be decoded to map the perturbation identity, typically iBAR 2, given the superior detection performance. Second and optionally, the first read can be stripped and the other iBAR, typically iBAR1, can be sequenced for as few as two additional cycles to filter out >93% of recombination events (**Methods**).

The use of iBARs and multiplexed decoding with CSM substantially improve decoding efficiency for optical pooled screens. During *in situ* sequencing, hands-on time, imaging duration, and sequencing reagent costs all scale linearly with the number of sequencing cycles required to decode library members. Decoding via spacer sequences, the standard approach for most CRISPR screens, including most optical pooled screens, typically requires sequencing all 20 bases for genome-scale libraries (**Figure 4G**). Imposing orthogonality requirements at the guide-design stage is one approach we have employed to improve decoding efficiency in CRISPR-KO screens, however this necessitates the selection of guides with compromised on- and off-target activities and is infeasible for applications with strict guide design constraints, including CRISPRa, CRISPRi, and tiling screens. The use of even a single linked barcode (e.g. an iBAR) can reduce the required cycle number for a given library size to roughly half that of decoding with the spacer sequence of a typical sgRNA library (**Figure 4G**). Serial detection of a second barcode can eliminate >93% of recombination events with as few as two additional cycles (**Methods**). This is analogous to the decoding strategy we suggest for T7-IVT approaches. With multiplexed detection of two linked barcodes (e.g. iBARs 1 and 2), the required number of cycles is again halved relative to a single linked barcode, as two bases are decoded per sequencing cycle (**Figure 4G**). For most libraries, a single additional sequencing cycle with multiplexed decoding is sufficient to detect >93% of recombination events, corresponding to a roughly 3-fold decrease in cycle number relative to decoding with spacer sequences (**Figure 4G, Methods**). In the current implementation, we anticipate this multiplexed decoding strategy is most appropriate for mRNA detection protocols.

Altogether, these approaches provide multiple options to efficiently decode optical pooled screens. For example, fully decoding a genome-wide CROPseq library with 4 guides per gene (about 80,000 constructs) would require sequencing all 20 cycles of the spacer. With CSM, an equivalent number of constructs, encoding twice as many guides per gene, could be decoded in as few as 10 cycles of sequencing for T7-IVT detection or 6 cycles of sequencing for mRNA detection, including recombination detection for both methods (**Figure 4G**).

## Discussion

CSM is a generalized multiplexing solution for single-target and combinatorial screens that enables Cas9-based perturbations, including CRISPR-KO and CRISPRi, and provides compatibility with high-content, single-cell phenotypic readouts including optical pooled screens and scRNA-seq screens. CSM addresses numerous technical challenges, including robust guide activity, low lentiviral recombination, and mRNA-barcoding. For optical pooled screens, CSM improves detection and decoding efficiencies, achieves superior single-target perturbation performance, and offers unique compatibility with combinatorial screens. CSMv2 retains all functionalities of the original CSM vector while adding compatibility for approaches using T7-IVT.

In single-target CRISPR-KO, CRISPRa, and CRISPRi screens, multiplexing systems typically achieve superior on-target performance with smaller library sizes compared to single-perturbation systems^12,13,15^. Our CRISPR-KO and CRISPRi viability screens with CSM support this claim (**Figure 2E-G, Supplementary Figure 5F-I**). While our viability screen results suggest there are marginal performance improvements beyond a single dual-guide construct per gene for CRISPR-KO and CRISPRi (**Figure 2E, F**), other screening approaches may have different requirements. Though dual-guide vectors offer improved performance with smaller libraries, selecting an optimal number of constructs per gene in a given screening context will still require balancing library size, coverage, and false positive and negative rates. We also demonstrate the applicability of CSM to combinatorial screens with as few as two constructs per gene or gene pair (six total for A, B, and AB) (**Figure 2K**). Beyond screening modalities commonly employed today, we expect CSM to be readily extensible to prime editing approaches^46,62^, which pair a prime editing gRNA (pegRNA) and nicking sgRNA or two pegRNAs.

It may be possible to extend CSM to three or more sgRNA perturbations, though challenges exist relating to molecular cloning and lentiviral performance. Commercially available oligo library synthesis options currently constrain length to about 300 nucleotides. Pending improvements in oligo synthesis, serial assembly steps may enable pooled cloning of higher order perturbation combinations. Both intermolecular and intramolecular lentiviral recombination are also expected to pose a greater challenge with extended arrays of repetitive elements; the incorporation of additional orthogonal sgRNA scaffolds and tRNAs to combat this effect will require careful testing and validation. Lentiviral titer may also be negatively impacted by encoding additional perturbations within the 3’ LTR, although we did not observe a difference in lentiviral titer between CSM constructs containing one or two sgRNAs (**Supplementary Figure 3B**). For these reasons, we expect Cas12a systems will remain attractive solutions for multiplexing 3 or more perturbations. We expect CSM-inspired designs (i.e. 3’ LTR-embedded antisense crRNA arrays) will enable multiplexed Cas12a screens with mRNA-barcoding compatibility.

For optical pooled screens, further improvements in decoding efficiency may be possible through a multiplexed detection approach; multiplexed readout of three or more barcodes would further reduce the requisite number of sequencing cycles to identify perturbations by decoding more bits of information per cycle. However, further increasing barcode readout multiplexity may be less impactful as additional reductions in sequencing reagent cost will be comparatively small and because phenotypic measurements at relatively high magnification may become the primary throughput bottleneck in the context of increasingly efficient library decoding.

Beyond decoding efficiency and detection of recombination events, multiple barcodes can enable additional functionalities such as barcoding of arbitrary subpopulations, including internal replicates and clonal populations. Barcoding of subpopulations in pooled CRISPR screens has been leveraged to improve enrichment screen performance^47,63,64^ and may be extensible to profiling screening techniques such as optical pooled screens and scRNA-seq-based screens. Additionally, clonal barcoding in screens may enable characterization of clonal heterogeneity and subclonal dynamics.

In conclusion, CSM is a versatile multiplexing platform for diverse CRISPR screening methodologies. We addressed challenges in multiplexing perturbations while maintaining design compatibility across enrichment, single-cell sequencing, and optical pooled screens. This versatility provides the opportunity for direct comparison and integration of different screening modalities, for example scRNA-seq and imaging-based approaches. CSM will enable single-target screens with smaller libraries and improved performance and we expect the compatibility of CSM with high-content screening techniques to enable new directions for combinatorial screens. In particular, we anticipate combinatorial optical pooled screens will be a powerful approach to interrogate genetic interactions at high throughput and with rich, single-cell-resolved phenotypic measurements.

## Supporting information

example vector maps

## Methods

### Arrayed molecular cloning

The CSM-Puro entry vector was derived from CROPseq-puro-v2 (Plasmid #127458) by modifying the lentiviral promoter and 3’ LTR with isothermal assembly. Alternate selection markers were subcloned from CSM-Puro by isothermal assembly.

CSM vectors are available on Addgene. To facilitate the generation of custom derivatives (e.g. alternative pol II promoters, selection genes, etc.), CSM vectors are supported in the Broad Institute Genetic Perturbation Platform’s modular vector assembly system, Fragmid^49^ (https://portals.broadinstitute.org/gppx/fragmid/public).

For arrayed cloning of CSM dual-guide vectors, dual-guide inserts were purchased as gene fragments from Twist Biosciences and cloned into the entry vector using golden gate assembly. Briefly, 10 nanograms (ng) of entry vector and 2 ng of gene fragment were assembled in a 2.5 µL reaction of 1X T4 DNA ligase buffer (New England Biolabs B0202S) and 1X BsmBI-v2 Golden Gate Enzyme Mix (New England Biolabs E1602S) with the thermal cycling protocol: 30 cycles of (42°C for 1 min, 16°C for 1 min), then 60°C for 5 min. All arrayed cloning was performed in NEB Stable chemically competent cells (New England Biolabs C3040H) according to the manufacturer’s recommendations and all cultures were grown at 30 °C. Liquid cultures were grown shaking at 225 RPM and 30 °C in 2xYT media with 100 µg/mL of carbenicillin.

### Pooled molecular cloning of CSM dual-guide libraries

For pooled cloning of CSM dual-guide vectors, 300 nucleotide oligo libraries were purchased from Twist Biosciences and cloned into the entry vector by restriction digestion and ligation. Code for the design of CSM oligo libraries is available on GitHub. Oligo pool amplification PCRs were assembled with 1X KAPA HiFi HotStart Ready Mix (Roche KK2601), 1X EvaGreen qPCR dye (Biotium 31000), 1M Betaine (MilliporeSigma B0300), 300 nM forward and reverse primers (Integrated DNA Technologies), and 12 pg/µL of template in 50 µL reactions. If multiple sublibaries with distinct amplification primer pairs are encoded in one oligo pool, the template concentration for the sublibrary is given by:

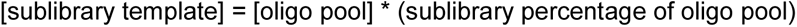

PCRs were conducted with the following thermal cycling protocol: 95 °C for 3 min, 14-16 cycles of (98 °C for 20 s, 62 °C for 15 s, 72 °C for 15 s), then 72 °C for 1 min. We strongly caution <u>against</u> the use of the manufacturer’s recommended amplification conditions, 10 ng of oligo pool template per 25 µL PCR (400 pg/µL), as of this publication date. Across three oligo pool orders, we observed that the frequency of recombination within plasmid libraries was determined by the oligo pool template concentration during amplification and template concentrations greater than 200 pg/µL resulted in double-digit recombination rates in plasmid libraries (**Supplementary Figure 4C**). For oligo pools that encode multiple sublibraries, the sublibrary concentration, not the total oligo pool concentration, determines recombination frequencies, so long as the amplification primers are specific to the sublibrary. Rather than high oligo pool template concentrations, we found scaling the number of amplification reactions was effective to maintain uniformity without compromising recombination.

Maintaining library uniformity requires amplifying a sufficient quantity of the oligo pool. While we found the manufacturer’s total ssDNA quantification is accurate, the amplifiable proportion of ssDNA (presumably the fraction of full-length products) varies as much as 20-fold between oligo pool orders when quantified by qPCR (**Supplementary Figure 4D**). Correspondingly, the ng amount of oligo pool needed to maintain library uniformity varies by the same factor. Pending manufacturer improvements in oligo pool synthesis, quantification, and/or quality control, we recommend performing qPCR to determine the concentration of amplifiable products. For small libraries, it may be most straightforward to skip oligo pool qPCR quantification and proceed with a conservative estimate of one 25 µL PCR reaction per 100 library members. For fewer PCR reactions, higher oligo template concentrations can be used but will result in predictably higher recombination rates (**Supplementary Figure 4C**).

For larger libraries, qPCR can be used to determine a minimum number of reactions necessary to maintain uniformity. Perform qPCR quantification of oligo pools alongside a standard curve to determine the concentration of full length oligos. A single oligo design purchased as a 300 bp gene fragment can serve as an appropriate standard. For our libraries, a per-oligo representation of 10^5^ in oligo pool amplification has been sufficient to maintain 90:10 ratios (the ratio in abundance of the 90th percentile construct to the 10th percentile construct) of about 2. Perform the requisite number of reactions to amplify 5x10^4^ full-length oligos per library member, while maintaining the effective template concentration to 12 pg/µL. Given the variability we have observed in oligo pool quantification, the minimum number of reactions for roughly 1,000-member libraries has varied between 1 and 15 reactions.

Following amplification, reactions were pooled and purified with the QIAquick PCR & Gel Cleanup Kit (Qiagen 28506) and quantified with the Qubit High Sensitivity dsDNA Quantitation assay (Thermo Fisher Scientific Q32854). Plasmid libraries were then constructed by restriction digestion and ligation. Entry vectors were digested with BsmBI-v2 (New England Biolabs R0739L), dephosphorylated with rSAP (New England Biolabs M0371L), and gel purified with the QIAquick PCR & Gel Cleanup Kit (Qiagen 28506). Amplified oligo libraries were digested with BsmBI-v2 (New England Biolabs R0739L) and purified with the QIAquick PCR & Gel Cleanup Kit (Qiagen 28506). The digested entry vector and amplified oligo libraries were quantified with the Qubit Broad Range dsDNA Quantitation assay (Thermo Fisher Scientific Q32853). Each 20 µL ligation reaction consisted of 20 femtomole (fmol) of BsmBI-digested amplified oligo pool, 20 fmol of digested backbone, 1X T4 DNA ligase buffer (New England Biolabs M0202S), and 20 U/µL of T4 DNA ligase (New England Biolabs M0202S). We observed that either 0.5:1 or 1:1 molar ratios of insert:backbone were optimal for library uniformity. Excess insert relative to backbone resulted in poor uniformity, with a clear bias determined by the identity of the ligation-adjacent base of iBAR 2 (**Supplementary Figure 4F**). Ligation reactions were incubated at 16 °C overnight, then heat-inactivated for 10 minutes at 65 °C. Reactions were purified with a 1.5X ratio of Ampure XP paramagnetic beads (Beckman Coulter A63880) and eluted in 10 µL of water. Purified ligations were electrotransformed by combining 5 µL of ligation and 25 µL of Endura electrocompetent cells (Biosearch Technologies 60242-2) in a 0.1 cm Gene Pulser cuvette (Biorad 1652083) on ice, electroporating on a Gene Pulser Xcell (Biorad 1652662) with the settings 1.8 kV, 600 Ohms, and 10 µF, and recovering for 90 minutes at 30 °C in 1 mL of Endura recovery media (Biosearch Technologies 60242-2). Cultures were then grown for 16 hours in 50 mL of 2xYT media with 100 µg/mL of carbenicillin, shaking at 225 RPM at 30°C. Plasmid libraries were then purified with a Plasmid Plus Midi Kit (Qiagen 12943) following the manufacturer’s instructions. We observed assembly and transformation efficiencies averaging about 300,000 colony forming units per fmol of digested backbone input.

CSM plasmid libraries were prepared for next-generation sequencing using a single-step PCR protocol to minimize amplification bias and PCR template switching. Plasmid libraries were amplified using 1X NEBNext Ultra II Q5 Master Mix (New England Biolabs M0544), 1X EvaGreen dye (Biotium 31000), 500 nM forward and reverse primers (Integrated DNA Technologies), and 100 ng of plasmid template per 25 µL reaction with the following thermal cycling conditions: 98 °C for 1 min, 10 cycles of (98 °C for 10 s, 67 °C for 10 s, 72 °C for 15 s), and 72 °C for 1 min. Reactions were purified with a 1X ratio of Ampure XP paramagnetic beads (Beckman Coulter A63880). To assay deletion recombination events (i.e. **Figure 1E**), we used primers flanking both sgRNA scaffolds and sequenced on a 300-cycle sequencing kit (e.g. Illumina MS-102-2002). For subsequent library sequencing without quantification of deletion events, we opted for primers that flank only the spacers and iBARs and used custom Illumina read primers (**Supplementary Table 2**). While this strategy does not capture the majority of deletion events, it enables the observation of all variable sequence elements (spacers, iBARs, and the middle tRNA) with 150-cycle sequencing kits (e.g. Illumina MS-102-3001) and positions spacers and iBARs as early as possible within NGS reads, where sequencing error rates are lowest. Code for CSM library NGS analysis is available on GitHub.

### Cell culture, lentiviral production, and transduction

HEK-293FT (Thermo Fisher Scientific R70007), doxycycline-inducible-Cas9 A549^65^ (a gift from J.T. Neal), wild-type A549 (ATCC), and doxycycline-inducible-Cas9 HeLa^16^ cell lines were maintained in DMEM(1X) + GlutaMAX (Gibco 10569010) supplemented with 10% (v/v) heat-inactivated fetal bovine serum (MilliporeSigma F4135), 100 U/mL Penicillin and 100 µg/mL Streptomycin (Gibco 15140122). hTERT-immortalized RPE1 cells with Zim3-dCas9-2A-BFP^24^ (a gift from Jonathan Weissman) were cultured in DMEM/F-12 + HEPES (Thermo Fisher Scientific 11330032) supplemented with 10% (v/v) heat-inactivated fetal bovine serum (MilliporeSigma F4135), 100 U/mL Penicillin and 100 µg/mL Streptomycin (Gibco 15140122), 0.01 mg/mL hygromycin B (Thermo Fisher Scientific 10687010). All cell lines were passaged with TrypLE (Thermo Fisher Scientific 12604013).

For lentiviral production, 6-well plates were seeded with 1 million HEK-293FT cells in 2 mL of media per well 24 hours prior to transfection. Lentiviral plasmids MD2.G (Addgene #12259), psPAX (Addgene #12260), and transfer plasmids were transfected at a mass ratio of 2:3:4 with Lipofectamine 3000 (Thermo Fisher Scientific L3000001) following the manufacturer’s protocol. The media was replaced 4 hours post-transfection. Lentivirus was harvested 48 hours post-transfection. Media was collected and centrifuged at 400 xg for 5 min at 20 °C, then the supernatant was collected and filtered with a 0.45 µM syringe-filter (VWR 28143-312). For “all-in-one” CSM vectors, expressing both Cas9 and the dual guides, we concentrated lentivirus 100-fold with Lenti-X Concentrator (Takara 631232). Lentivirus was then stored at -80 °C in single-use aliquots.

Functional lentiviral titers were determined by transduction in the appropriate cell line with a lentiviral dilution series and quantifying survival. For A549 and HeLa cells, lentiviral transduction was performed by mixing cells in media supplemented with 8 µg/mL of polybrene (Santa Cruz Biotechnology 28728-55-4) with lentivirus and centrifuging in cell culture plates at 1000 xg and 33 °C for 2 hours. Media was replaced 24 hours post-transduction. Puromycin selection was initiated with the addition of 1 µg/mL puromycin for A549 cells or 2 µg/mL puromycin for HeLa cells (Thermo Fisher Scientific A1113803) 24 hours post-transduction. Cell Titer Glo (Promega G7573) was used to quantify survival after 48 hours of Puromycin selection, comparing survival against minus-virus/minus-selection positive control and minus-virus/plus-selection negative control. For RPE1 cells, lentiviral transduction was performed by adding lentivirus and 8 µg/mL of polybrene (MilliporeSigma TR-1003) to cells 24-hours after seeding. Media was replaced 24 hours post-transduction with 10 µg/mL puromycin (Thermo Fisher Scientific A1113803). Cell Titer Glo (Promega G7573) was used to quantify survival 4 days post-transduction. Cells were passaged in-well once to accelerate puromycin selection as the RPE1 line is resistant to low puromycin concentrations.

Percent survival was converted to MOI assuming a Poisson distribution for transduction:

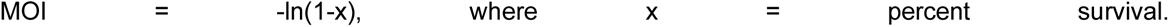

Dilutions of virus yielding 10%-60% survival were used to calculate titers as high and low extremes are more sensitive to experimental noise. Viral titer (genomic integrations per µL virus) was then determined from MOI:

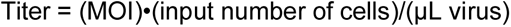

### Pooled viability screens

For the knockout screens, we selected all guides using CRISPick^8,9^. For the CRISPRi screen, we selected two guides from a published screen^24^ and four guides using CRISPick^8,9^. All steps of the pooled viability screens were performed to maintain library representation of at least 1000X across at least two independent biological replicates. For the A549 CRISPR-KO screen, wild-type A549 cells were transduced with lentivirus at a MOI of 0.1 and selected with 1 µg/mL puromycin starting 24 hours post-transduction. We harvested cells at 14 days post-transduction. For the HeLa CRISPR-KO screen, doxycycline-inducible Cas9 HeLa cells were transduced with lentivirus at a MOI of 0.1 and selected with 2 µg/mL puromycin for four days, starting 24 hours post-transduction. To initiate Cas9 knockout, doxycycline was added at 2 µg/mL and cells were harvested at 14 days post-transduction. For the RPE1 CRISPRi screen, RPE1 cells expressing Zim3-dCas9-P2A-BFP were transduced with lentivirus at a MOI of 0.1. We added 10 µg/mL puromycin (Thermo Fisher Scientific A1113803) 24-hours post-transduction and harvested cells at days 7 and 14 post-transduction.

Genomic DNA was purified with the QIAamp DNA Blood Midi Kit (Qiagen 51183). Libraries were then prepared for NGS using a single-step PCR. Genomic DNA libraries were amplified using 1X NEBNext Ultra II Q5 Master Mix (New England Biolabs M0544), 1X EvaGreen dye (Biotium 31000), 500 nM forward and reverse primers (Integrated DNA Technologies), and up to 2.5 µg of genomic DNA template per 50 µL reaction with the following thermal cycling conditions: 98 °C for 1 min, 24 cycles of (98 °C for 10 s, 67 °C for 10 s, 72 °C for 15 s), and 72 °C for 1 min. Optionally, lower concentrations of gDNA can be used to improve amplification efficiency, though it is important to scale the number of reactions to maintain sufficient library coverage. Reactions were pooled and purified with the QIAquick PCR & Gel Cleanup Kit (Qiagen 28506). We then gel purified reactions with the QIAquick PCR & Gel Cleanup Kit (Qiagen 28506). Libraries were quantified with the Qubit High Sensitivity dsDNA Quantitation assay (Thermo Fisher Scientific Q32854) and sequenced on an Illumina MiSeq with a 150-cycle kit (Illumina MS-102-3001) with 75 cycles for each read and 8 cycles for each index. Code for NGS analysis is available on GitHub.

### Quantification of endogenous editing by next-generation sequencing

Doxycycline-inducible-Cas9 A549 cells were transduced and selected with puromycin for five days as described above. To induce Cas9 expression and editing, transduced cells were cultured for 7 days with 1 µg/mL of doxycycline. Genomic DNA was harvested by discarding culture media and adding lysis buffer consisting of 20 mM Tris pH 8 and 0.1% Triton X-100 (MilliporeSigma T9284) with 60 ng/mL of Proteinase K (New England Biolabs P8107S) added immediately prior to use. Lysate was incubated at 65 °C for 6 min, then 95 °C for 2 min, and stored at -20°C. Endogenous loci were amplified with 1X Q5 High-Fidelity Master Mix (New England Biolabs M0492), 1 M Betaine (MilliporeSigma B0300), 500 nM forward and reverse primers (Integrated DNA Technologies), and gDNA in cell lysate of at least 5,000 cells, and the following thermal cycling protocol: 98 °C for 1 min, then 30 cycles of (98 °C for 10 s, 62 °C for 10 s, and 72 °C for 20 s), then 72 °C for 2 min. Amplicons were then barcoded for NGS by combining 2 µL of amplicon from the previous PCR with 1X Q5 High-Fidelity Master Mix (New England Biolabs M0492) and 500 nM forward and reverse indexing primers (Integrated DNA Technologies), and cycling with the following conditions: 98 °C for 1 min, then 10 cycles of (98 °C for 10 s, 60 °C for 10 s, and 72 °C for 15 s), then 72 °C for 2 min. Amplicons were then pooled, gel purified with the QIAquick PCR & Gel Cleanup Kit (Qiagen 28506), quantified with the Qubit High Sensitivity dsDNA Quantitation assay (Thermo Fisher Scientific Q32854), and sequenced on an Illumina MiSeq. Genome editing was quantified with CRISPResso2^66^.

### Direct-capture single-cell RNA sequencing with 10X Genomics 5’ capture

The direct-capture protocol for CSM follows the Chromium GEM-X Single Cell 5’ Reagent Kits v3 with Feature Barcode technology for CRISPR Screening (10X Genomics CG000735), with the following modifications that substitute primers and amplification conditions that are appropriate for the design of CSM. The protocol was evaluated in HeLa cells.

The reverse transcription mastermix was prepared by combining the RT Reagent E (10X Genomics 2001106), Poly-dT RT Primer B (10X Genomics 2001110), Reducing Agent B (10X Genomics 2000087), RT Enzyme E (10X Genomics 2001105/2001146) – 27.5 µL in total – and adding two custom reverse transcription primers: 0.5 µL of 10 µM sgRNA 1 RT primer (oRW1220) and 0.5 µL of 10 µM sgRNA 2 RT primer (oRW1221). The sgRNA scaffold modifications require the use of these custom RT primers. The RT primer included in the CRISPR Poly-dT Primer Mix B (10X Genomics 2001145) does not have perfect complementarity to both sgRNAs.

Following GEM generation, reverse transcription, and cDNA amplification, cDNA was purified via SPRI-select. During the SPRI-select cDNA cleanup, the cDNA library was size-separated into smaller cDNAs, for the sgRNA library, and larger cDNAs, for the gene expression library, as previously described^13^ and documented in commercial protocols (e.g. 10X Genomics CG000735). Following purification, the larger fraction was processed for gene expression library construction following the standard protocol (10X Genomics CG000733). Following purification, sgRNAs were amplified from the smaller cDNA fraction with the following PCR mix: 5 µL of small-fraction cDNA template, 500 nM of each primer, and 1X NEBNext Ultra II Q5 Master Mix (New England Biolabs M0544) in a 100 µL reaction. Three primers were used: a universal forward primer (oRW1223) and one reverse primer for each sgRNA (oRW1226 for sgRNA 1 and oRW1227 for sgRNA 2). Both sgRNAs are amplified in the same reaction. The PCR was run with the following program: 98 °C for 1 min, then 11 cycles of (98 °C for 10 s, 67 °C for 10 s, and 72 °C for 15 s), then 72 °C for 1 min. sgRNA amplicons were purified with a 1X ratio SPRIselect and eluted in 30 µL of 10 mM Tris-Cl, pH 8.5. sgRNA amplicons were then prepared for NGS with a sample-indexing PCR, purified, and quantified as described in the commercial protocol (10X Genomics CG000735).

Gene expression and sgRNA libraries were pooled at a 10:1 molar ratio and sequenced on an Illumina NovaSeq X with 28 cycles for read 1, 10 cycles for each index, and 90 cycles for read 2. Cellranger was used to assign reads and sgRNAs to cells using a custom alignment reference for sgRNAs.

### In situ amplification - mRNA protocol

The mRNA *in situ* amplification protocol was modified from our previous studies^16,61^. Cells were fixed in 4% (v/v) formaldehyde (Electron Microscopy Sciences 15714) and 0.007% (v/v) glutaraldehyde (Electron Microscopy Sciences 16120) in 1X PBS (Ambion AM9625) for 30 minutes at room temperature, then washed twice in PBS. The use of glutaraldehyde in the primary fixation step can impact some immunofluorescence stains; omission or titration to lower concentrations may offer a balance of detection sensitivity and compatibility with phenotype measurements. Samples were permeabilized in 1X PBS + 0.2% Tween-20 (VWR 100216-360) for 15 minutes at room temperature, then washed twice in 1X PBS + 0.1% Tween-20, henceforth “PBS-T”. For phenotypic measurements prior to reverse transcription and cDNA-fixation, we recommend using RiboLock Rnase Inhibitor (Thermo Fisher Scientific EO0384) together with RNase-free reagents to preserve mRNA integrity.

The reverse transcription solution was prepared with the following composition: 1X RevertAid RT buffer (Thermo Fisher Scientific EP0452), 250 µM dNTPs (New England Biolabs N0447L), 1 µM each biotinylated reverse transcription primer (Integrated DNA Technologies), 200 µg/mL molecular biology grade recombinant albumin (rAlbumin) (New England Biolabs B9200S), 0.8 U/µL RiboLock RNase inhibitor (Thermo Fisher Scientific EO0384), and 4.8 U/µL RevertAid H minus Reverse Transcriptase (Thermo Fisher Scientific EP0452). Samples were incubated in reverse transcription solution for 16 hours at 37 °C. Samples were then washed twice with PBS-T and incubated for 15 minutes with 20 µg/mL Streptavidin (New England Biolabs N7021S) and 100 µg/mL rAlbumin (New England Biolabs B9200S) in 1X PBS. Next, samples were washed twice with PBS-T prior to post-fixation in 3% formaldehyde (Electron Microscopy Sciences 15714) and 0.1% glutaraldehyde (Electron Microscopy Sciences 16120) for 30 minutes at room temperature. After fixation, samples were washed twice with PBS-T and incubated in gapfill and ligation solution at 37 °C for 5 min, followed by 45 °C for 90 minutes. The gapfill and ligation solution was composed of 1X Ampligase buffer (Lucigen A3210K), 50 nM dNTPs (New England Biolabs N0447L), 0.1 µM each padlock probe, 200 µg/mL rAlbumin (New England Biolabs B9200S), 0.4 U/µL RNase H (Enzymatics Y9220L), 0.02 U/µL TaqIT polymerase (Enzymatics P7620L), and 0.5 U/µL Ampligase (Lucigen A3210K). Samples were then washed twice with PBS-T and incubated in RCA solution for 16 hours at 30 °C. RCA solution was composed of 1X Phi29 buffer (Thermo Fisher Scientific EP0091), 5% glycerol (MilliporeSigma G5516), 250 µM dNTPs (New England Biolabs N0447L), 200 µg/mL rAlbumin (New England Biolabs B9200S), and 1 U/µL Phi29 DNA polymerase (Thermo Fisher Scientific EP0091). Following RCA, samples were washed twice in PBS-T and incubated with 1 µM each sequencing primer (Integrated DNA Technologies) in 2X SSC buffer (Ambion AM9763) for 30 minutes at room temperature, followed by two PBS-T washes.

### In situ amplification - T7 IVT protocol

The T7-IVT *in situ* amplification protocol was modified from previous studies^4,5^. Unless noted otherwise, samples were prepared with a PFA fixation protocol as follows. Cells were fixed in 4% (v/v) formaldehyde (Electron Microscopy Sciences 15714) in 1X PBS (Ambion AM9625) for 30 minutes at room temperature, then washed twice in PBS. Cells were permeabilized for 30 minutes in 70% ethanol, followed by three washes with PBS-T. Cells were decrosslinked via incubation in 0.1M sodium bicarbonate and 0.3 M NaCl in water at 65 °C for 4 hours, followed by three PBS-T washes. As noted in previous work^4^, it is critical to use plates that tolerate high temperatures (e.g. Cellviz, P96-1.5H-N).

The IVT solution was prepared as follows: 1X T7 Reaction Buffer (NEB E2040L), 10 mM each NTPs (NEB E2040L), 5 mM DTT (NEB E2040L), 0.4 U/µL RiboLock RNase inhibitor (Thermo Fisher Scientific EO0384), and 1X to 0.2X diluted T7 RNA Polymerase Mix (NEB E2040L - M0255AVIAL). To reduce cost, we found dilutions of the T7 RNA Polymerase Mix down to 0.2X (relative to the manufacturer’s recommendations) produced comparable detection efficiencies for CSMv2. For dilution of the T7 RNA Polymerase Mix, additional NTPs can be purchased separately (NEB N0466L) and T7 Reaction Buffer can be purchased individually by special order (NEB B2041AVIAL). Unless noted otherwise, all data was generated with 0.2X T7 RNA Polymerase Mix. IVT reactions were incubated at 37 °C overnight (12-16 hours).

After IVT, cells were washed three times with PBS-T and then fixed for 30 minutes at room temperature in 3% formaldehyde (Electron Microscopy Sciences 15714) and 0.1% glutaraldehyde (Electron Microscopy Sciences 16120). Fixation was quenched by incubating for 1 minute at room temperature in 0.2 N Tris-HCl pH 8, followed by three washes with PBS-T. Subsequently, cells were processed as described in the mRNA *in situ* amplification protocol, as described above, beginning with the addition of the reverse transcription solution.

For samples prepared with methanol/acetic acid fixation, cells were fixed in 3:1 methanol:acetic acid for 20 minutes at room temperature. After three PBS-T washes, samples were processed as described above, starting from incubation with the IVT solution.

### Sequencing by synthesis

Sequencing by synthesis was then performed as previously described^16,61^. Briefly, samples were incubated in incorporation mix (MiSeq Nano kit v2 reagent 1) (Illumina MS-103-1003) for 5 minutes at 60 °C on a flat-top thermal cycler, then washed six times with PR2 buffer (Illumina MS-103-1003), followed by 5 heated washes in PR2, 5 min each at 60 °C. Samples were imaged in 2X SSC + 200 ng/mL DAPI (MilliporeSigma D9542) on a Nikon Ti2 Microscope at 10X magnification. To proceed to the next cycle, samples were incubated in cleavage mix (MiSeq Nano kit v2 reagent 4) (Illumina MS-103-1003) for 6 min at 60 °C, then washed three times in PR2 followed by three heated PR2 washes of 1 min each at 60 °C. Samples were then ready to return to the incorporation step for the subsequent sequencing cycle. To strip primers for serial decoding approaches, samples were washed twice in 2xSSC and 80% formamide at 60 °C for 5 minutes, followed by three washes with PBS-T. Then the next sequencing primer was hybridized, as described above. *In situ* sequencing images were analyzed as previously described^16,61,67^.

### Comparison of decoding efficiencies of *in situ* detection approaches

To determine cycling requirements for spacer sequences, we performed 50 simulations at each library size by randomly sampling guides from the Dolcetto genome-wide CRISPRi library^9^. The same approach applied to sgRNAs sampled from CRISPR-KO (Brunello) and CRISPRa (Calabrese) libraries (data not shown) were indistinguishable from the CRISPRi library. Single barcode encoding was approximated by the step function:

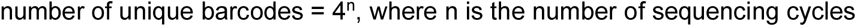

For dual barcodes, we assumed that reads could be assigned to the appropriate iBAR position in a sequence-independent manner, such as through the use of a fluorophore-conjugated oligo to label one iBAR. Dual barcode encoding was approximated by the step function:

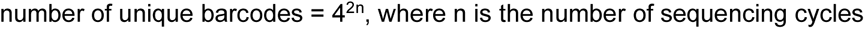

Dual barcode encoding with 95% recombination detection was approximated by the step function:

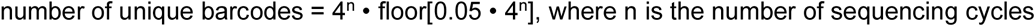

Practically, barcodes will be constrained by restriction enzyme sites, homopolymers, and GC content. Barcodes may be restricted further for edit distance to enable error detection and/or error correction, however we have not explored these considerations in this simple approximation.

In our library design pipeline, we have implemented user-specified edit distance guarantees and recombination detection simulation. In most library designs with diverse iBAR sequences, achieving >93% recombination detection typically requires adding just one extra cycle for multiplexed decoding (for 2 additional bases total) or two additional cycles for serial decoding. Intuitively, this reflects the probability of a recombination event resulting in the same two additional bases that correspond to the unrecombined design, which is (¼)^2^ or 6.25%, rendering the remaining 93.75% of events detectable recombination. With this level of filtering, a library with 10% recombination, would yield a filtered population >99% recombination-free.

## Acknowledgements

We would like to thank the following individuals for helpful discussions and feedback on the manuscript: Michael Ward, Andrew Bassett, John Doench, Christoph Bock, Rebecca Carlson, Mohamad Najia, Frances Keer, as well as all members of the Blainey Lab. Andrew Bassett engaged throughout the project in helpful discussions about molecular cloning workflows and tRNA-based sgRNA multiplexing systems. John Doench helped make reagents available through the Broad Institute Genetic Perturbation Platform’s modular vector assembly platform, Fragmid. We thank Manuel Lessi, Nicolò Caporale, Giuseppe Testa, and Natan Pirete for their expertise with single-cell RNA sequencing workflows. We would like to acknowledge Rebecca Dertinger for administrative support and Robert Majovski for manuscript proofreading. R.T.W. is supported by the National Science Foundation Graduate Research Fellowship Program under Grant No. 1745302. Y.Q. is supported by the National Institutes of Health (NIH) under grant K00 CA264422 and by funding from the Eric and Wendy Schmidt Center at the Broad Institute of MIT and Harvard. Byunguk K is supported by NIH R01 NS 089076, NIH 1R01 MH128366, and Simons Foundation Autism Research Initiative award #890477. We acknowledge the in-kind contribution of Chromium reagents from 10X Genomics used in this work.

## Author contributions

R.T.W. conceived of the project, designed and performed experiments, analyzed data, and wrote the manuscript. Y.Q., Bryce K., J.O.A., M.T., and Byunguk K, designed and performed experiments, analyzed data, and edited the manuscript. P.C.B. supervised the work and edited the manuscript.

## Data availability

Data is available upon request.

## Code availability

Code is available on GitHub:

- CSM library design and sequencing analysis: https://github.com/rtwalton/CROPseq-multi
- DNA barcode design: https://github.com/feldman4/dna-barcodes
- *in situ* sequencing data analysis: https://github.com/feldman4/OpticalPooledScreens, https://github.com/cheeseman-lab/brieflow

## Conflict of Interest

R.T.W., Y.Q., and P.C.B. are inventors on patents relating to optical pooled screening technologies. PCB is a consultant to and/or holds equity in companies that develop or apply biotechnologies: 10X Genomics (whose products - both purchased and provided in-kind - were used in this work), General Automation Lab Technologies/Isolation Bio, Celsius Therapeutics, Next Gen Diagnostics, Cache DNA, Concerto Biosciences, Stately, Ramona Optics, Bifrost Biosystems, and Amber Bio. His laboratory has received research funding from Calico Life Sciences, Merck, and Genentech for work related to genetic screening.

## Supplementary Figures

**Supplementary Figure 1.**
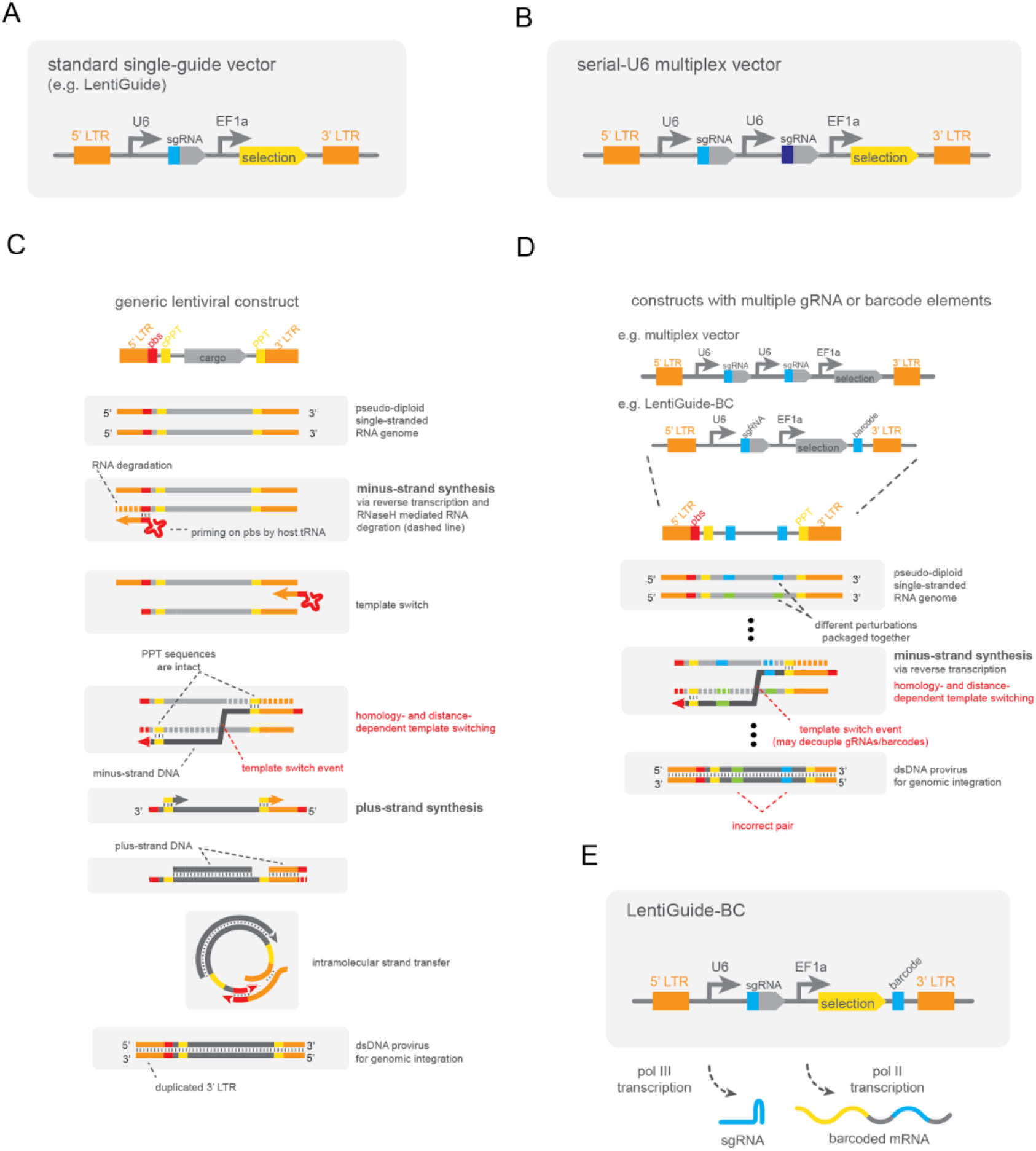
Lentiviral vectors for CRISPR perturbation. (A-B) Schematic representations of a standard single-guide lentiviral vector (A) and derivative multiplexing systems (B). (C-D) Processing of the lentiviral RNA genome into double stranded DNA for genome integration for generic vectors (C) and those with multiple guides or barcode elements (D) to illustrate steps vulnerable to recombination. One template switch is shown for simplicity, however multiple template switches are expected given a length of several kilobases switching rate of roughly 1 per 0.5-1 kilobases^50^. Illustration inspired by Adamson *et al*.^27^ (E) Schematic representations of LentiGuideBC vector design for pairing guide RNAs with mRNA barcodes. LTR, long terminal repeats; pbs, primer binding site; PPT, polypurine tract; cPPT, central polypurine tract.

**Supplementary Figure 2.**
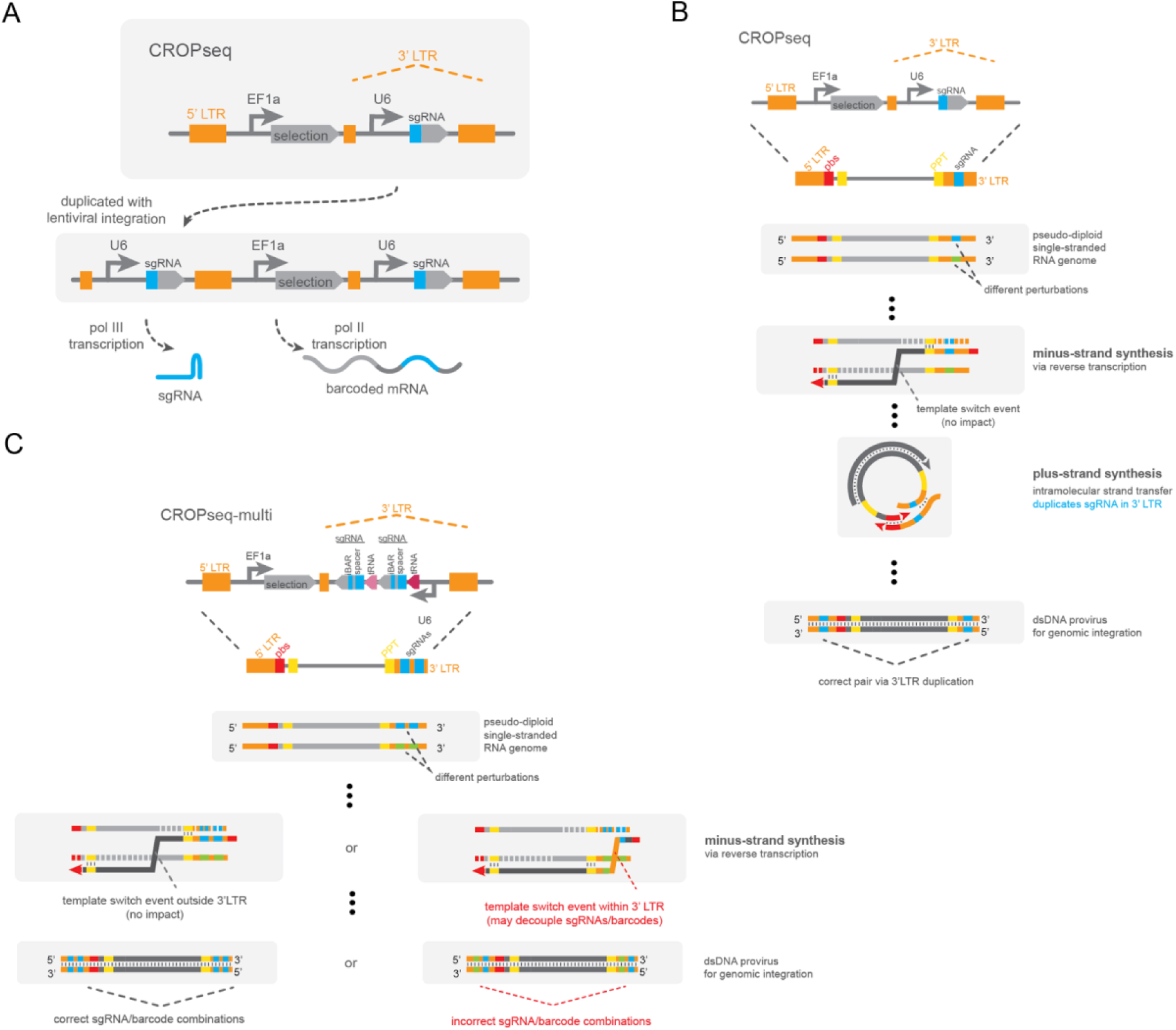
CROPseq and CROPseq-multi lentiviral vectors. (A) Schematic representation of the CROPseq vector design. (B) Abbreviated schematic of lentiviral RNA genome processing into double stranded DNA for genome integration for CROPseq vectors, highlighting the intramolecular 3’ LTR duplication that is not vulnerable to recombination. (C) Abbreviated schematic of lentiviral RNA genome processing into double stranded DNA for genome integration for CROPseq-multi vectors. CROPseq-multi remains vulnerable to recombination during minus strand synthesis, but is robust against recombination during 3’ LTR duplication. LTR, long terminal repeats; pbs, primer binding site; PPT, polypurine tract; cPPT, central polypurine tract.

**Supplementary Figure 3.**
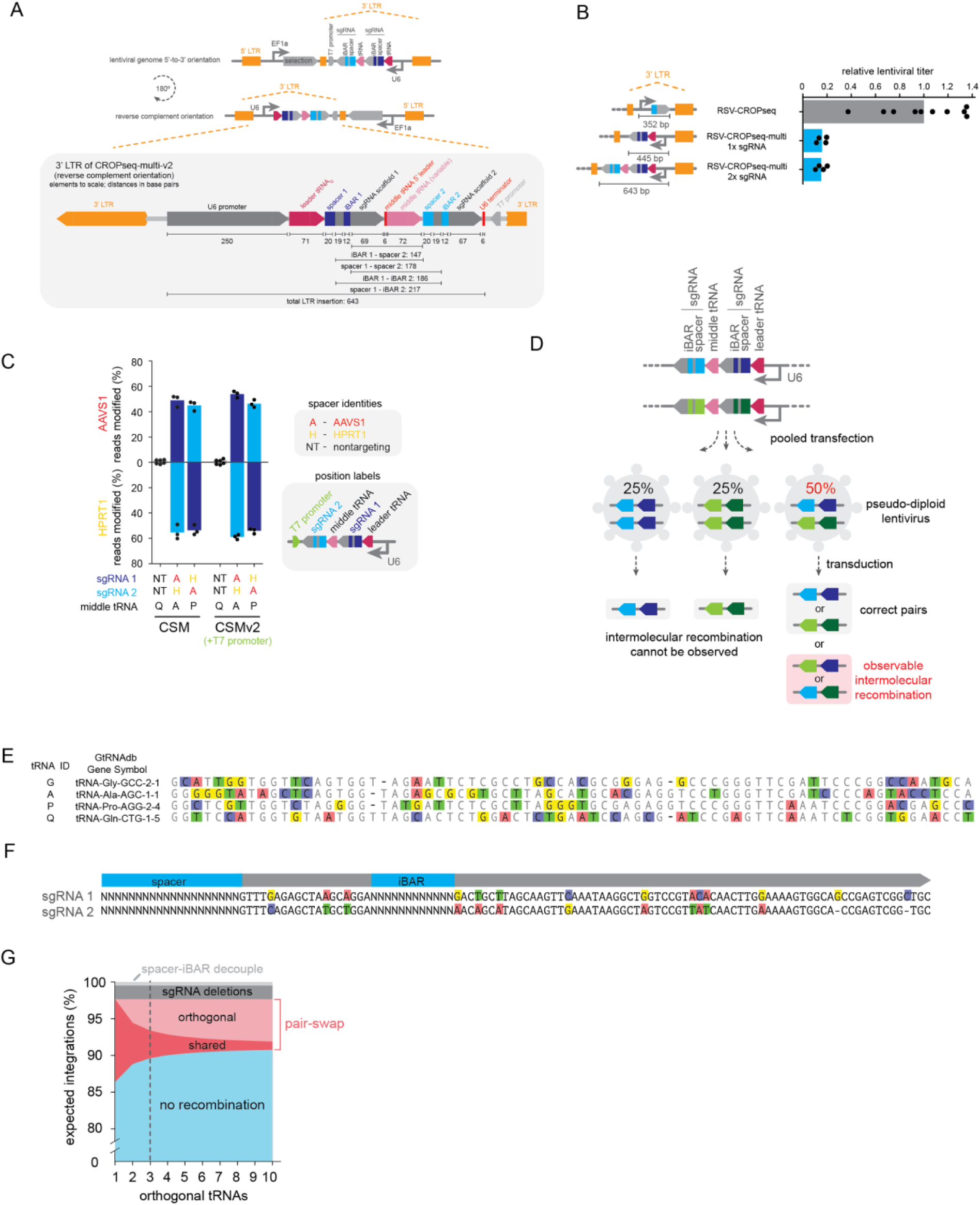
Detailed molecular design and additional performance evaluation of CROPseq-multi. (A) Detailed illustration of the CROPseq-multi-v2 3’ LTR design. (B) Lentiviral titers of RSV-CROPseq-multi vectors relative to CROPseq. (C) Genome editing activities of CROPseq-multi and CROPseq-multi-v2 vectors in SpCas9-expressing A549 lung adenocarcinoma cells, quantified by next-generation sequencing. Mean and n=3 biological replicates shown. (D) Assaying lentiviral recombination with pooled lentiviral production. (E) Sequence alignment of orthogonal tRNAs employed in CROPseq-multi. (F) Sequence alignment of orthogonal sgRNA scaffolds used in CROPseq-multi. (G) Expected recombination frequencies in CROPseq-multi libraries with orthogonal tRNAs. Expected recombination rates for a three-tRNA system indicated with a dotted line. LTR, long terminal repeats; RSV, Rous sarcoma virus.

**Supplementary Figure 4.**
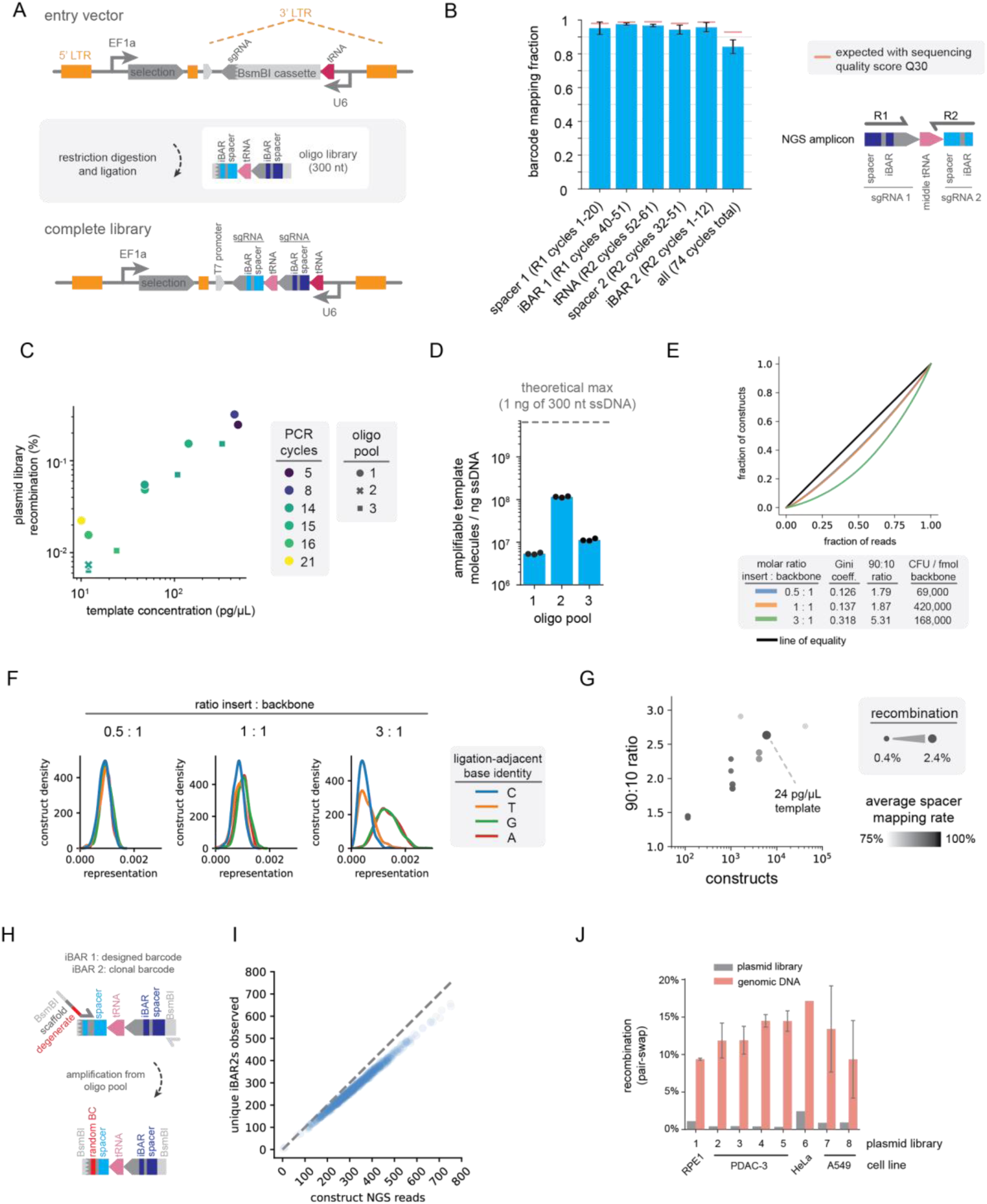
Construction and quality control of CROPseq-multi libraries. (A) Schematic of pooled CROPseq-multi library construction. (B) Mapping rates of individual barcode elements in a CROPseq-multi library, determined by next-generation sequencing. “All” is the mapping rate of all elements (spacer 1, iBAR1, 10 bases of the middle tRNA, spacer 2, and iBAR 2) in a read. In addition to oligo synthesis and amplification errors, sequencing error rates explain a fraction of unmapped barcode elements. (C) Recombination in CROPseq-multi plasmid libraries as a function of effective template concentration and polymerase chain reaction amplification cycle number. (D) Polymerase chain reaction quantification of three oligo pool orders. (E) Lorenz plots, Gini coefficients, and 90:10 ratios for 1,080-member CROPseq-multi plasmid libraries built with different assembly conditions. 90:10 ratios are the ratio in abundance of the 90th percentile construct to the 10th percentile construct. (F) Restriction-ligation assembly bias based on the identity of the variable ligation-adjacent base (encoded by iBAR2) is corrected by equimolar or limiting ratios of insert to backbone. (G) Summary metrics for additional CROPseq-multi libraries constructed with optimized conditions. Unless noted otherwise, libraries were constructed with optimal amplification conditions (12 pg/µL template). (H) Schematic of CROPseq-multi oligo library amplification for clonal barcoding applications. (I) High diversity of iBAR2 sequences for a 1,080-member, clonally-barcoded CROPseq-multi library exceeds next-generation sequencing depth. (J) Recombination with CROPseq-multi across distinct plasmid and cell libraries. Mean and standard deviation shown. LTR, long terminal repeats; NGS, next-generation sequencing; ssDNA, single-stranded DNA

**Supplementary Figure 5.**
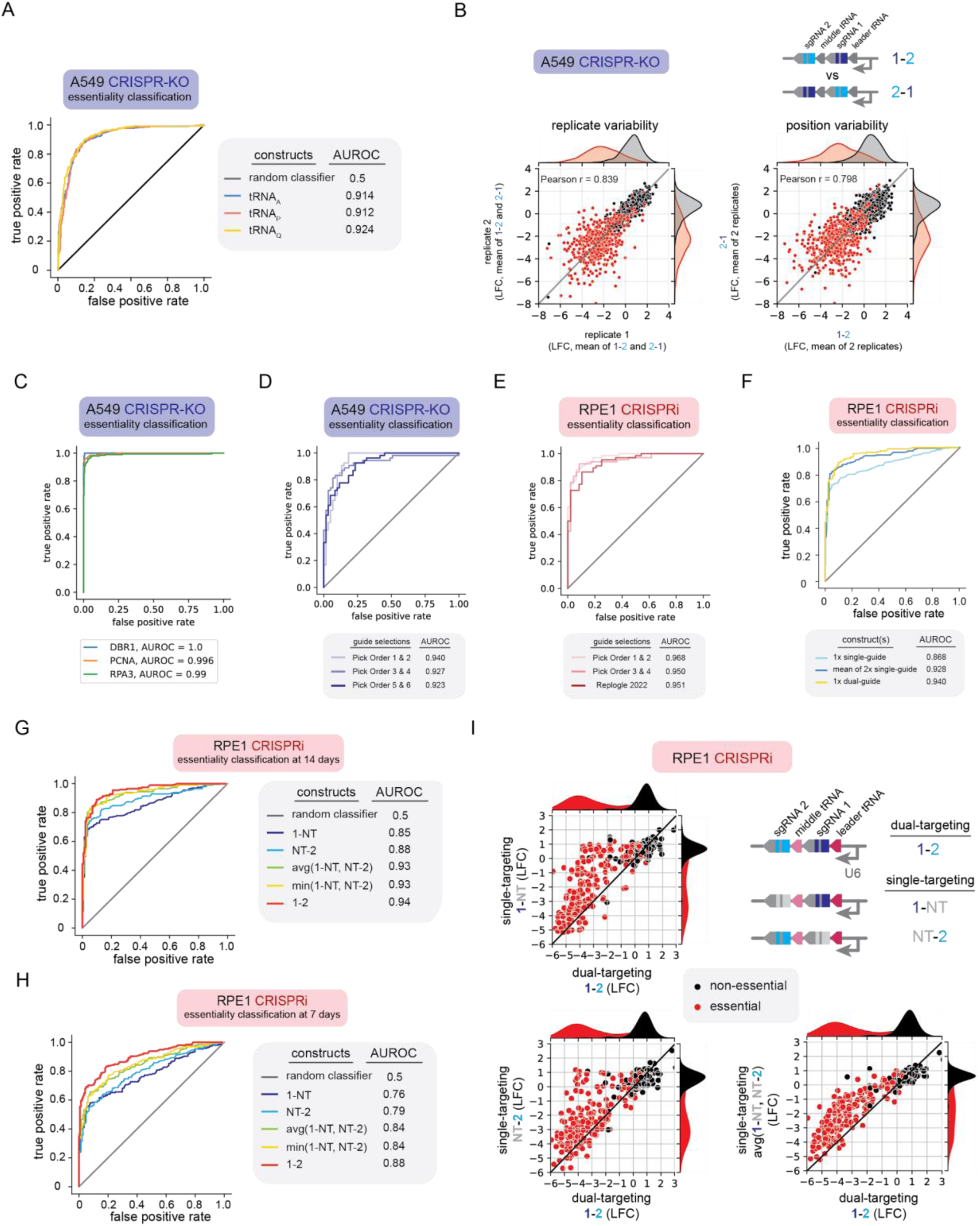
Additional benchmarking of CROPseq-multi in pooled viability screens. (A) Gene essentiality classification for identical sets of dual-guide constructs, varying only by the identity of the middle tRNA. (B) Scatterplots showing the correlation of guide pairs across biological replicates, averaged across two constructs with matched guide pairs in opposite orientations (left, Pearson r = 0.839) and the correlation of constructs with matched guide pairs in opposite orientations, averaged across two biological replicates (right, Pearson r = 0.798) (C) Gene essentiality classification for individual constructs targeting essential genes DBR1, PCNA, and RPA3, varying only by 12 nucleotide iBAR sequences. (D, E) Gene essentiality classification for dual guide constructs by *a priori* computational ranking of guide pairs for both CRISPR-KO (D) and CRIPSRi (E) screens. (F) Essentiality classification for CRISPRi in RPE1 cells at day 14 for individual single-guide constructs, the average of two single-guide constructs, and individual dual-guide constructs. (G,H) Essentiality classification for CRISPRi in RPE1 cells at days 14 (G) and 7 (H) for individual single-guide constructs, including dual-targeting, single-targeting, and composite metrics. (I) Log2 fold changes of dual-targeting (1-2) CROPseq-multi constructs versus the corresponding single-targeting (1-NT and NT-2) constructs and construct averages. Log2 fold changes are the mean of at least two biological replicates. LFC, log2 fold change; NT, non-targeting; AUROC, area under receiver operator characteristic curve.

**Supplementary Figure 6.**
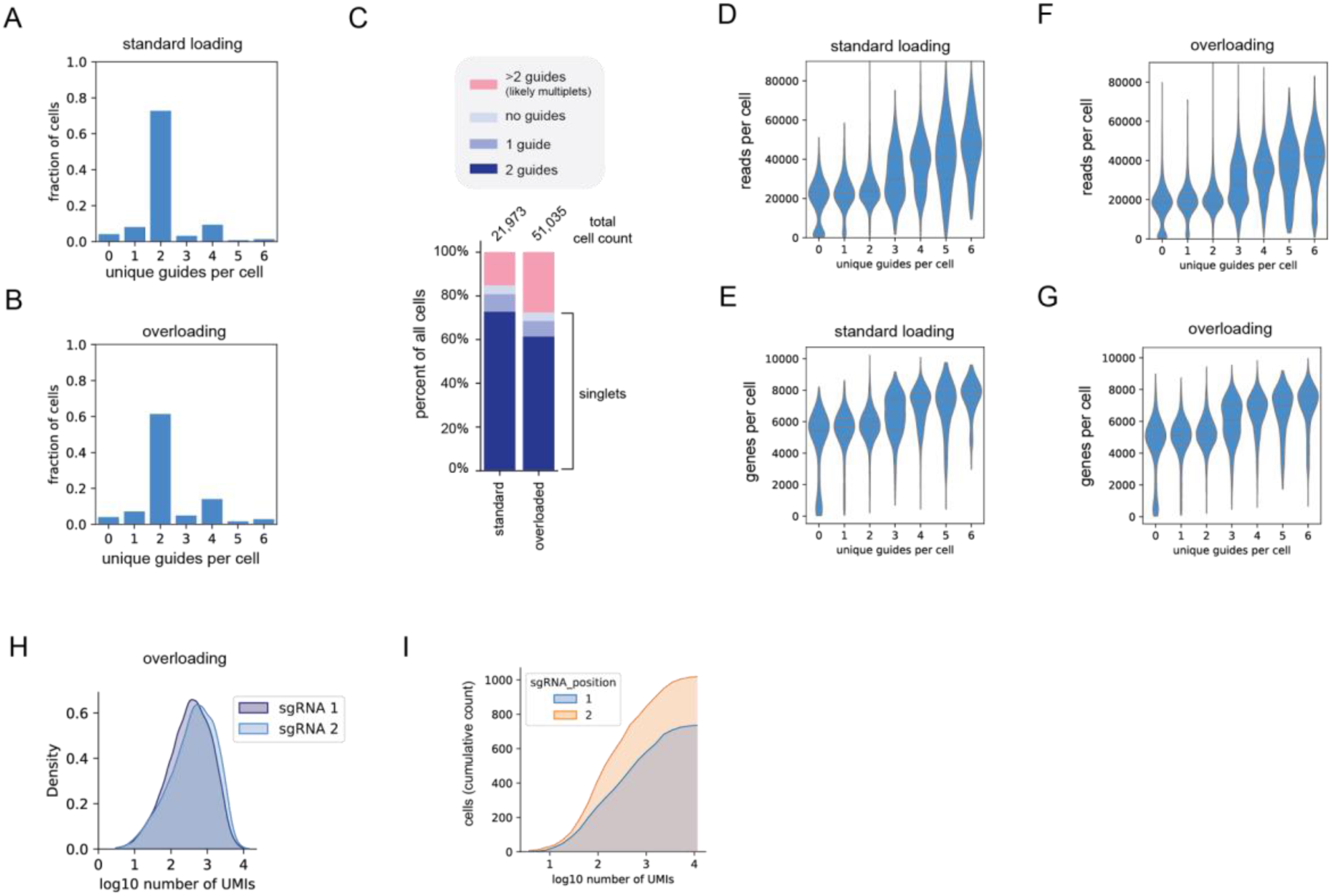
Performance metrics for CROPseq-multi detection with 5’ direct-capture single-cell RNA-sequencing. (A-B) The number of unique guides assigned to cells for standard (A) and overloading (B) conditions in HeLa cells. (C) Percentage of all “cells” (cell barcodes identified with Cell Ranger) by the number of detected sgRNAs. (D-E) The number of reads (D) and genes (E) detected per cell with standard cell loading conditions, groupbed by the number of unique guides detected per cell. (F-G) The number of reads (F) and genes (G) detected per cell with cell overloading conditions, groupbed by the number of unique guides detected per cell. (H) Kernel density estimate of unique molecular identifiers per sgRNA per cell for cells with exactly 2 sgRNAs detected with cell overloading conditions. Quartiles shown on violin plots. (I) Cumulative cell counts for unique molecular identifiers per sgRNA, by sgRNA position, for cells with only one observed sgRNA. UMI, unique molecular identifier.

**Supplementary Figure 7.**
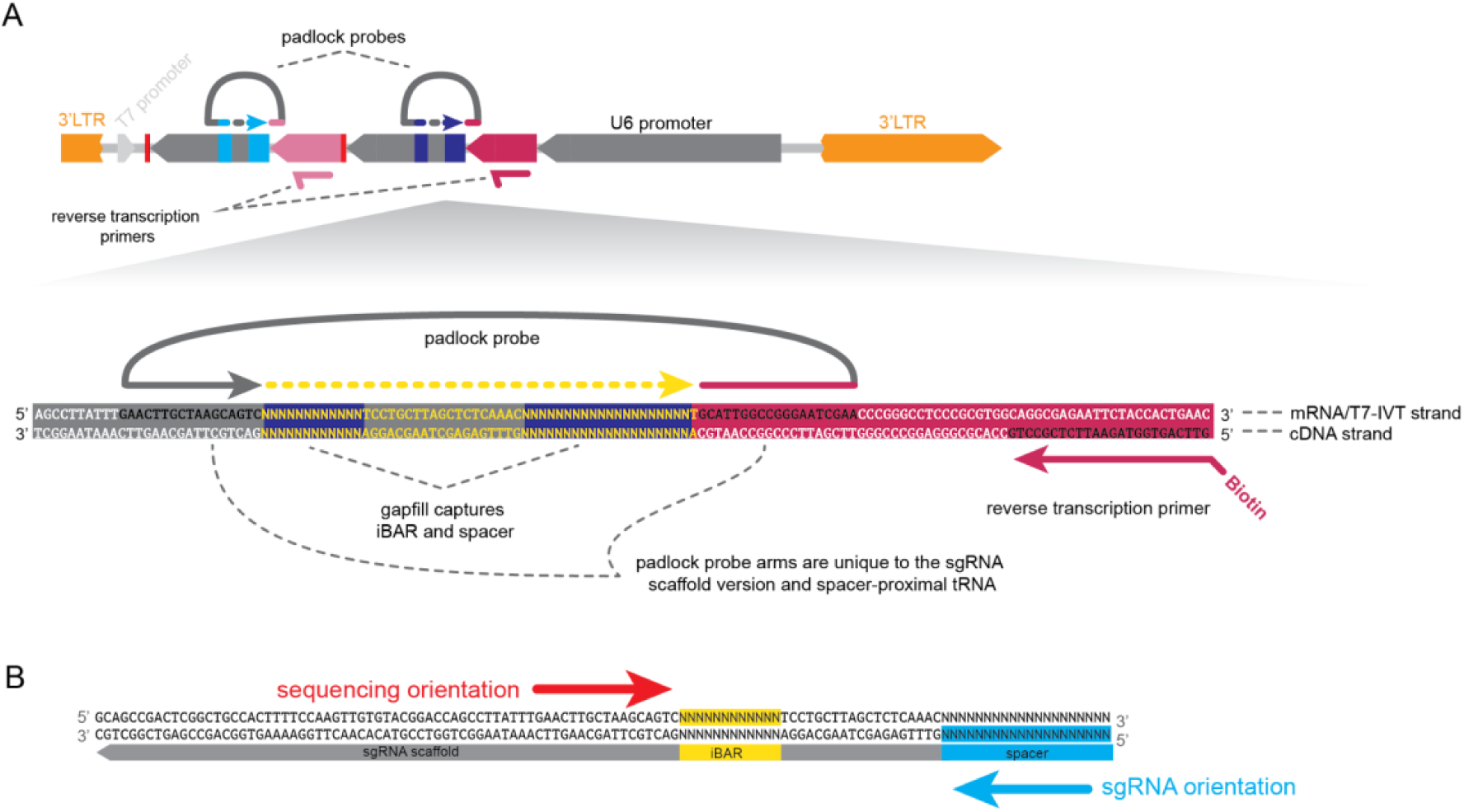
Detailed molecular design of CROPseq-multi *in situ* detection workflows. (A) Illustration of oligonucleotide reagent design for *in situ* detection of CROPseq-multi barcodes. CROPseq-multi-v2 is shown. (B) Illustration of the orientation of *in situ* sequencing relative to the sgRNA. LTR, long terminal repeats; IVT, *in vitro* transcription.

**Supplementary Figure 8.**
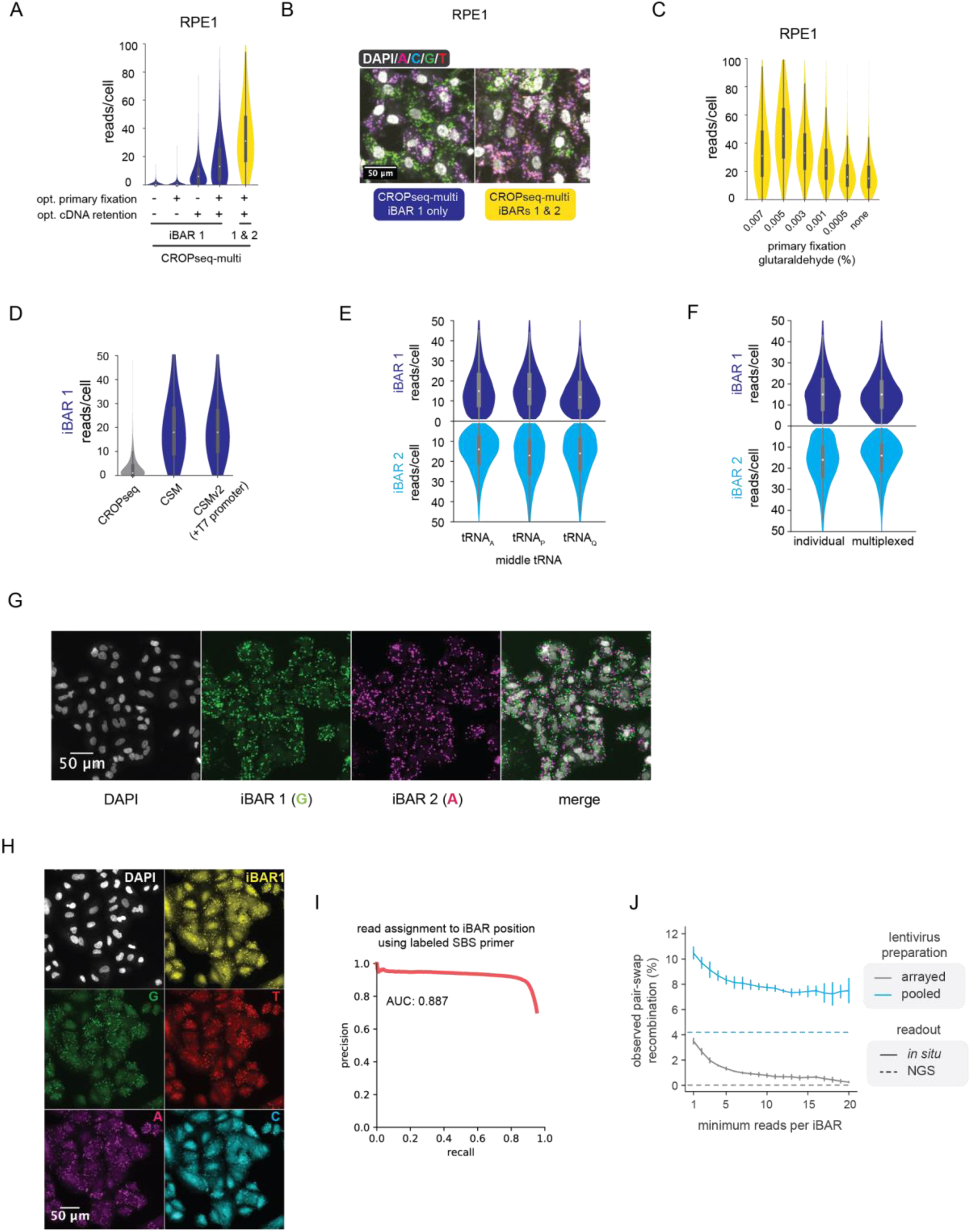
Additional validation of CROPseq-multi mRNA *in situ* detection. (A) Optimization of CROPseq-multi *in situ* mRNA detection in RPE1 cells. (B) Representative images of CROPseq-multi *in situ* mRNA detection in RPE1 cells with an optimized protocol. (C) Optimization of glutaraldehyde concentration in the primary fixation step for multiplexed *in situ* mRNA detection of CROPseq-multi in RPE1 cells. (D) Comparison of mRNA detection efficiency between CROPseq, CROPseq-multi, and CROPseq-multi-v2. (E) Comparison of the mRNA detection efficiency, in A549 cells, of iBARs 1 and 2 across constructs employing three orthogonal middle tRNAs. (F) mRNA detection efficiency of CROPseq-multi iBARs individually and with multiplexed detection. (G) Representative images of *in situ* sequencing reads with multiplexed detection in A549 cells. (H) Representative image showing the selective labeling of iBAR 1 reads with a fluorescent oligo in a multiplexed detection experiment, together with DAPI-stained nuclei and the four sequencing bases in separate fluorescent channels. (I) Precision-recall curve for assignment of individual reads to either iBAR 1 or iBAR 2 on signal from the iBAR-1-specific probe. (J) Quantification of recombination in a three-vector pool measured via *in situ* sequencing and NGS. AUC, area under curve; NGS, next generation sequencing.

**Supplementary Figure 9.**
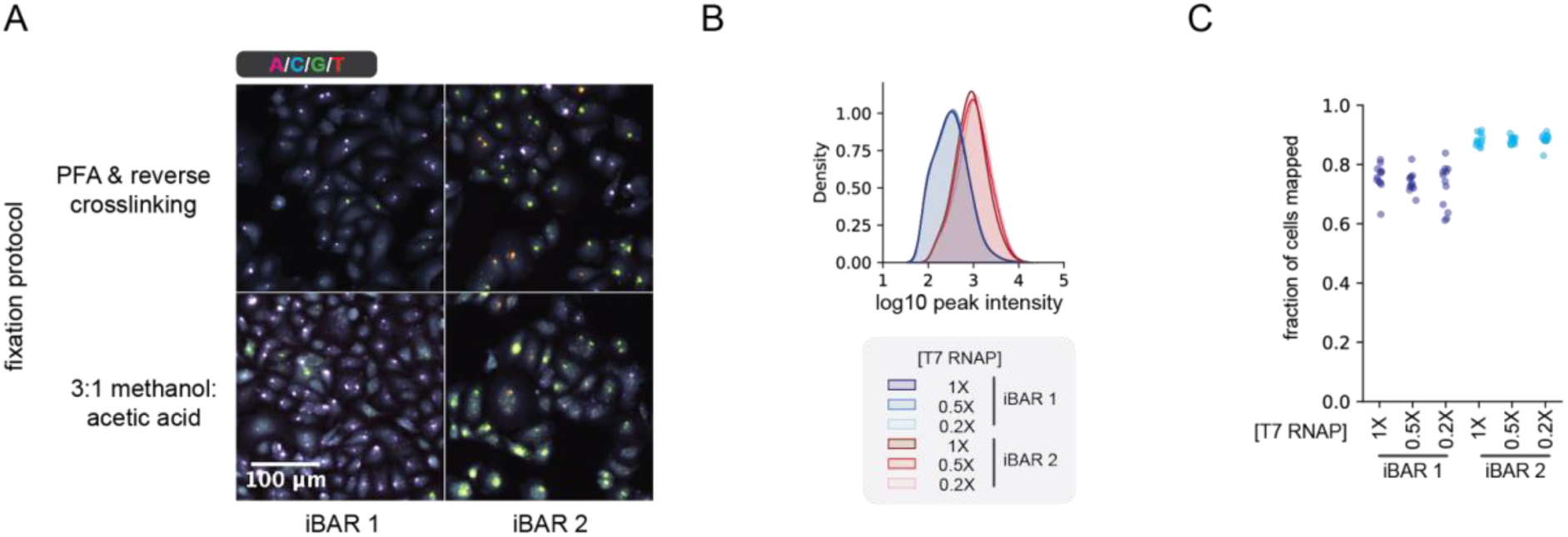
Additional validation of CROPseq-multi T7 *in vitro* transcription *in situ* detection. (A) Representative images of *in situ* detection of CROPseq-multi iBARs 1 and 2 with T7-IVT detection protocols. (B, C) Peak intensities (B) and fraction of cells mapped to the library (C) for T7-IVT detection of CROPseq-multi iBARs 1 and 2 with dilutions of T7 RNA polymerase. All conditions display at least three technical replicates and at least three fields of view per replicate. Unless stated otherwise, all T7-IVT detection results used a PFA fixation and decrosslinking protocol. PFA, paraformaldehyde; RNAP, RNA polymerase; IVT, *in vitro* transcription.

## Supplementary Data

**Supplementary Table 1.**
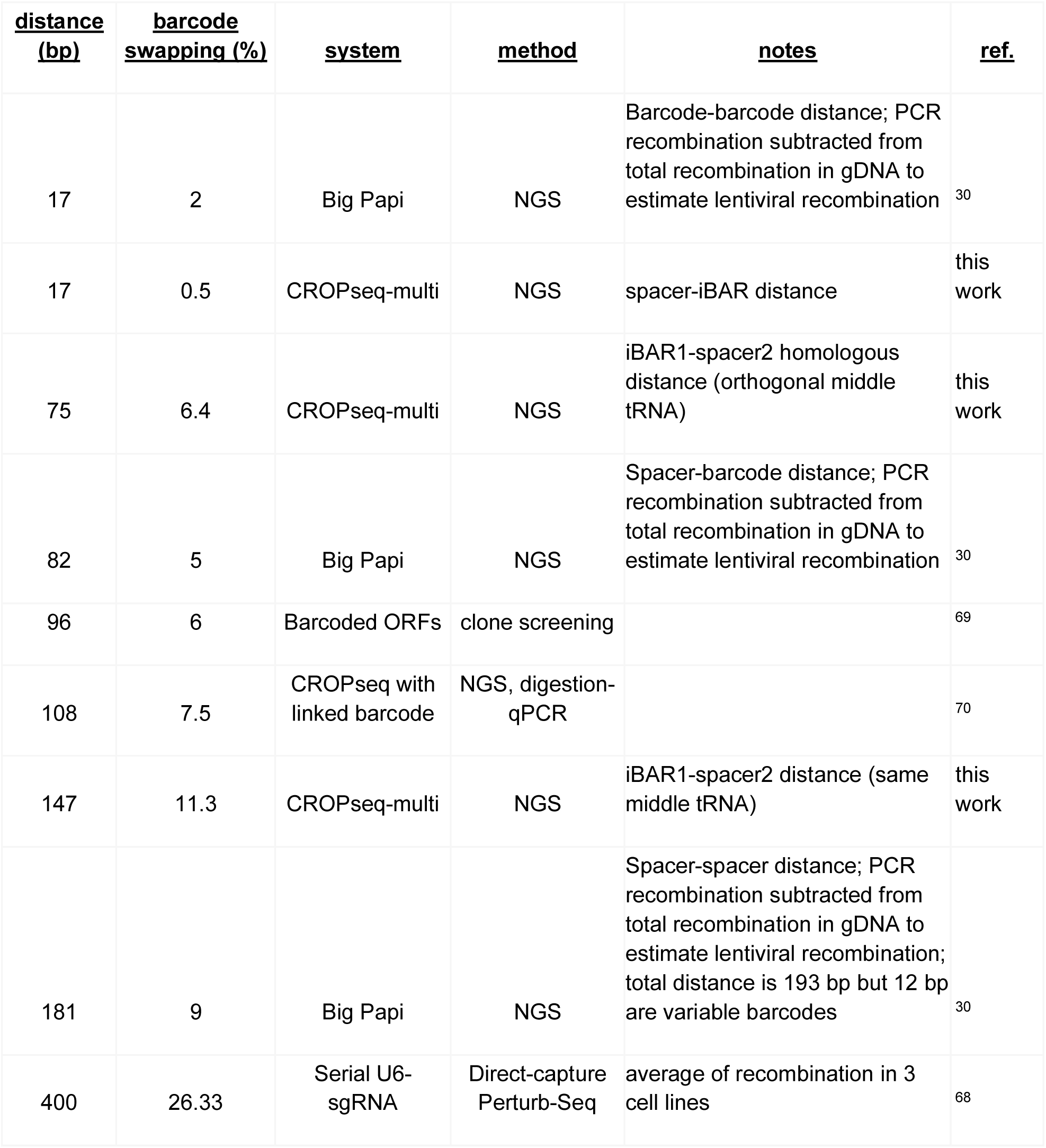
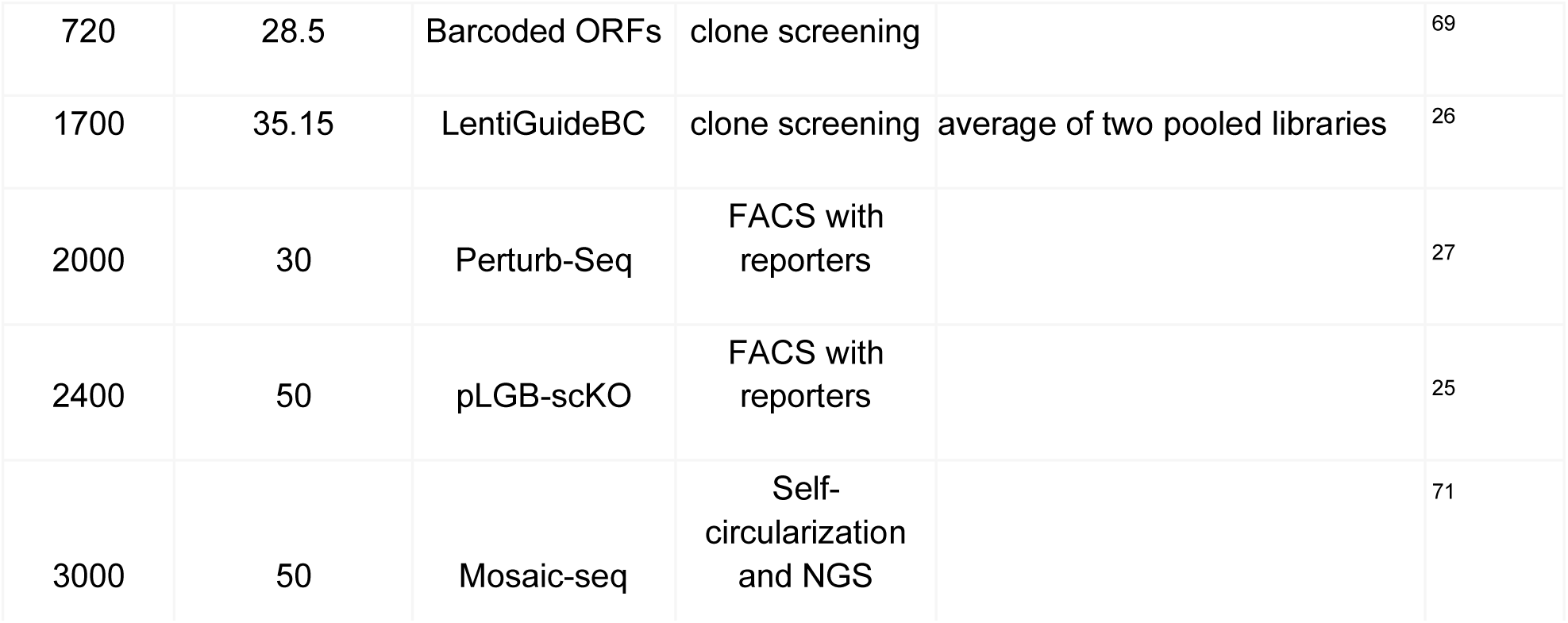
Reporter barcode swapping rates with lentiviral barcoding systems. NGS, next-generation sequencing; FACS, fluorescence activated cell sorting; qPCR, quantitative polymerase chain reaction; ORF, open reading frame.

**Supplementary Table 2** - see separate supplementary files

## Notes

### Summary of Updates

This update includes three major additions: (1) an updated design, CROPseq-multi-v2, that adds compatibility for T7 in vitro transcription-based barcode detection methods, (2) validated methodology for single-cell RNA-sequencing readouts, and (3) additional pooled screens validating the performance of CROPseq-multi for CRISPR-KO and CRIPSRi, as well as single-target and combinatorial screening applications.

